# Temporal gating dictates stress-induced transcript export from the nucleus

**DOI:** 10.64898/2026.04.14.718555

**Authors:** Noah S. Helton, Benjamin Dodd, Stephanie L. Moon

## Abstract

Prior studies have largely focused on transcriptional and translational control during stress, but how regulated nuclear mRNA export contributes to the stress response remains unresolved. We show that nuclear mRNA export is progressively inhibited during arsenite and heat stress in human cells. In contrast to previous work largely in yeast that suggests nuclear export of stress-induced transcripts is prioritized through sequence-specific mechanisms, we demonstrate that temporal gating determines the nucleocytoplasmic distribution of mRNAs during stress. Using single molecule mRNA imaging and transcriptome-wide analyses, we find the majority of stress-induced mRNAs, including heat shock protein transcripts, accumulate in the nucleus during stress. However, a subset of stress-induced mRNAs, notably *HMOX1, JUN*, and *FOS,* escape nuclear retention. mRNAs transcribed early during stress, including those encoding immediate early genes, redox mediators, and protein chaperones, are exported from the nucleus prior to the global inhibition of mRNA export. In contrast, mRNAs transcribed later are retained in the nucleus until stress is resolved. Reporter RNA assays confirm that transcriptional timing determines mRNA export competence. This work reveals that the timing of transcription, rather than transcript-specific sequence features, is the major determinant of nuclear export efficiency of stress-induced transcripts in human cells.

## Introduction

Eukaryotic cells respond to stress by remodeling gene expression at the transcriptional and translational levels. Stress-induced genes are rapidly transcribed and many are translated via upstream open reading frames while the majority of constitutively expressed transcripts are translationally repressed (Pakos-Zebrucka et al. 2016). The heat shock response (HSR) and the integrated stress response (ISR) coordinate gene expression upon exposure to stressful conditions including environmental changes, toxins, or in disease contexts (Pakos-Zebrucka et al. 2016; Lindquist 1986; Harding et al. 2003; Morimoto 1998; Vihervaara et al. 2018). During the HSR, dissociation of the transcription factor HSF1 (heat-shock factor 1) from heat shock proteins (HSPs) drives transcription of chaperones critical for resolving misfolded, aggregation-prone proteins (Morimoto 1998; Abravaya et al. 1992; Baler et al. 1992; Shi et al. 1998). During the ISR, preferential translation of transcription factors including *ATF4* drives expression of stress-induced genes (Han et al. 2013; Young and Wek 2016; Vattem and Wek 2004; Harding et al. 2000). Both pathways are self-limiting through negative feedback loops. Increased levels of HSP70 leads to sequestration of HSF1 to repress the HSR (Morimoto 1998; Abravaya et al. 1992; Baler et al. 1992; Shi et al. 1998), and the stress-induced gene *GADD34* mediates P-eIF2α dephosphorylation to inhibit the ISR (Novoa et al. 2001; Connor et al. 2001; Novoa et al. 2003). Commonly used stressors that activate both the HSR and ISR in mammalian cells are arsenite and heat (typically 42 - 45°C) (Johnston et al. 1980; Li 1983; Frydrýšková et al. 2020; Taniuchi et al. 2016; Lu et al. 2001). While the mechanisms underlying transcriptional and translational control of stress-induced genes have been extensively characterized, how cells temporally coordinate stress-induced gene expression, and the role of regulated nuclear mRNA export in this process, remain unclear.

The export of mRNA from the nucleus is coupled to transcription and pre-mRNA processing through the sequential assembly and remodeling of messenger ribonucleoprotein (mRNP) complexes. Generally, mRNAs are co-transcriptionally licensed for export by the TREX complex (THO complex, UAP56, and Aly/Ref), which scaffolds interactions between mRNPs and the NXF1-NXT1 heterodimer to facilitate export to the cytoplasm through the nuclear pore complex (NPC) (Pühringer et al. 2020; Strässer and Hurt 2000, 2001; Katahira et al. 1999; Hautbergue et al. 2008; Masuda et al. 2005; Mor et al. 2010; Ben-Yishay et al. 2019; Viphakone et al. 2019; Müller-McNicoll et al. 2016; Luo and Reed 1999; Pacheco-Fiallos et al. 2023). Additionally, RNA modifications (Yang et al. 2017; Roundtree et al. 2017; Lee et al. 2026), mRNA transit through nuclear speckles (Wang et al. 2018; Khanna et al. 2014; Fan et al. 2018), and specialized export factors (Aksenova et al. 2020; Culjkovic et al. 2006; Culjkovic-Kraljacic et al. 2020; Mars et al. 2024; Lee et al. 2020) can play non-mutually exclusive roles in mRNA export. Misprocessed mRNAs are generally retained in the nucleus and subject to nuclear degradation (Hilleren et al. 2001; Rambout and Maquat 2024; Gordon et al. 2021; Tian et al. 2026b), while spliced mRNAs are rapidly exported (Luo and Reed 1999; Zhou et al. 2000; Valencia et al. 2008; Prasanth et al. 2005). Nuclear mRNA export is not generally thought to be a rate-limiting step or a major contributor to the regulation of gene expression. Rather, export is a quality control step that safeguards gene expression fidelity and is the default outcome immediately following pre-mRNA processing.

Despite historically being viewed as the default pathway for properly processed mRNAs, emerging evidence suggests that regulated nuclear export can shape gene expression. For example, recent kinetic modeling analyses suggest nuclear export should be considered as a rate-limiting step in gene expression (Steinbrecht et al. 2024; Müller et al. 2024). Additionally, mRNA retention in the nucleus has been proposed to mitigate noise in gene expression due to bursty transcription in mammalian cells (Bahar Halpern et al. 2015). Nuclear mRNA retention is also implicated in the neuronal response to stimuli and stress responses. Certain pre-mRNAs can accumulate in the nucleus and undergo induced splicing, bypassing the need for transcription to rapidly alter gene expression in neurons (Mauger et al. 2016; Mazille et al. 2022). Similarly, the pre-mRNA encoding the nutrient transporter mCAT2 is spliced upon inflammatory stress in mouse fibroblasts (Prasanth et al. 2005). Further, studies primarily done in baker’s yeast demonstrate bulk poly(A)+ RNA levels increase in the nucleus during stress, which may suggest global nuclear RNA export inhibition (Saavedra et al. 1996; Tani et al. 1995; Coban et al. 2024; Zander et al. 2016). However, RNA accumulation in the nucleus does not necessarily result from impaired nuclear export. RNA levels could increase in the nucleus if transcription rates are higher than nuclear export rates, if decay of unspliced or misprocessed mRNAs in the nucleus is inhibited, or if the cytoplasmic pool of mRNA degrades faster than nuclear mRNAs. Distinguishing between these possibilities requires time-resolved, quantitative analyses of nuclear mRNA export during stress.

How nuclear mRNA export contributes to stress-induced gene regulation in mammalian cells remains unresolved. While several studies have examined nucleocytoplasmic distribution of heat shock protein mRNAs and other stress-induced transcripts, there is no consensus mechanism that explains how their localization is regulated during stress. The preferential export of HSP mRNAs has been proposed to occur via promoter-and/or sequence- specific mechanisms, including direct binding to export factors or interaction with antisense RNAs (Saavedra et al. 1996; Coban et al. 2024; Zander et al. 2016; Skaggs et al. 2007a). Changes in splicing have also been proposed to contribute to differential nuclear mRNA export during heat stress in human and *Drosophila* cell cultures. Increased levels of unspliced HSP and other mRNAs occurs during heat stress, with G-rich motifs suggested to mediate sequence-specific licensing of mRNA export (Yost and Lindquist 1986; Bond 1988; Shalgi et al. 2014). Additionally, alternative export factors including eEF1A1, TPR, and THOC5/Aly, or transit through nuclear speckles, have been implicated in *HSP70* export during heat stress (Katahira et al. 2009, 2019; Vera et al. 2014; Hu et al. 2010; Khanna et al. 2014; Wang et al. 2018). Together, these studies propose multiple sequence- and factor- specific mechanisms for the nuclear export of specific mRNAs, yet whether these mechanisms are generalizable and explain the nucleocytoplasmic distribution of stress-induced mRNAs more broadly has not been established.

In this study, we demonstrate that nuclear export of mRNAs is progressively inhibited during arsenite and heat stress in human cells. This decline in nuclear export activity acts as a temporal gating mechanism that dictates which mRNAs are released into the cytoplasm during stress. Using transcriptional shutoff assays, single molecule mRNA imaging, and global analysis of nuclear and cytoplasmic RNAs, we find the majority of heat shock protein and other stress-induced gene mRNAs are retained in the nucleus during stress. We define a subset of 55 mRNAs including immediate early genes and protein chaperones that are induced during stress and are primarily localized in the cytoplasm, suggesting they escape nuclear retention. The nuclear retention of stress-induced transcripts is independent of splicing, as mRNAs lacking introns can be either retained in, or exported from, the nucleus. Transcriptome-wide time-course experiments demonstrate that early-transcribed stress-induced genes escape nuclear retention, while later-transcribed mRNAs remain within the nucleus. The export of all stress-induced gene mRNAs examined is progressively inhibited, regardless of whether they are transcribed early or late during stress. Inducible reporter mRNAs also accumulate in the nucleus over time during stress, and fail to export from the nucleus during late arsenite or heat stress. The inhibition of nuclear mRNA export is regulated and reversible, as stress-induced genes and induced reporter mRNAs rapidly accumulate in the cytoplasm during the recovery from arsenite or heat stress. Together, we demonstrate that mammalian cells regulate nuclear export to facilitate the selective export of early-transcribed genes while retaining most mRNAs within the nucleus until stress is resolved.

## Results

### HSP70 mRNAs accumulate in the nucleus during arsenite and heat stress

While analyzing the subcellular localization of stress-induced mRNAs (Helton et al. 2025), we serendipitously observed that the HSP70 mRNAs *HSPA1A* and *HSPA1B* accumulate to high levels in the nucleus in U-2 OS cells during arsenite stress. Arsenite is widely used to dissect stress response pathways as it is highly reproducible, easily controllable without changing environmental conditions, is a therapeutic and environmental stressor, causes oxidative stress, and robustly induces the ISR and HSR (Carlin et al. 2016; Paul et al. 2023; Helton et al. 2025; McEwen et al. 2005; Kedersha et al. 2000; Johnston et al. 1980; Li 1983; Frydrýšková et al. 2020). Our observation that *HSPA1A/B* accumulate in the nucleus suggests the nuclear export of these mRNAs is inhibited during stress. To begin to test this hypothesis, we first evaluated the nuclear and cytoplasmic localization of *HSPA1A* during a time course of stress and the recovery from stress. We treated U-2 OS cells with arsenite (250 µM) and fixed them at 45 or 105 min, and also evaluated cells recovering from stress (for 45 min) at 60 or 120 min following arsenite washout. The U-2 OS cells expressed endogenously tagged GFP-G3BP1 to demarcate the cytoplasm, and nuclei were stained with Hoechst. We performed single-molecule fluorescence *in situ* hybridization (smFISH) to determine the percentage of *HSPA1A* mRNAs in the nucleus during stress and recovery. We also assessed if bulk polyadenylated RNAs are globally increased in the nucleus during stress using oligo(dT) probes, which detect constitutive and stress-induced mRNAs. As controls, we used the cytoplasmic-enriched mRNA *GAPDH* and the nuclear-enriched lncRNA *NEAT1*. We applied a modified BigFISH pipeline (Imbert et al. 2022; Helton et al. 2025) to segment nuclei and cytoplasm and extract the number of RNAs from each compartment.

We observed that *HSPA1A* significantly accumulated (by ∼3-fold) in the nucleus at 45 and 105 min of arsenite stress, and returned to baseline by 120 min recovery from stress (**Figure 1A**). This phenomenon was not cell-type specific, as iPSC-derived motor neurons and immortalized retinal pigment epithelial hTERT-RPE cells also exhibited nuclear accumulation of *HSP70* mRNAs during arsenite stress (**Supplemental Fig. S1**). As expected, the levels of *HSPA1A* significantly increased during and after recovery from arsenite stress in U-2 OS cells (**Supplemental Fig. S2A**). Polyadenylated RNAs also slightly but significantly accumulated in the nucleus during stress, and returned to pre-stress levels during the recovery from stress (**Figure 1B**). The constitutively expressed *GAPDH* was enriched in the cytoplasm (∼83 - 87%) and lncRNA *NEAT1* was enriched in the nucleus (∼92 - 97%) during stress (**Figure 1C&D**). There were no significant changes in the abundance of poly(A)+ RNA, *GAPDH*, or *NEAT1* during or after stress (**Supplemental Fig. S2A-D**). Therefore, *HSPA1A* and, to a lesser extent, bulk polyadenylated RNA, accumulate in the nucleus during arsenite stress and appear to redistribute into the cytoplasm after stress.

**Figure 1.**
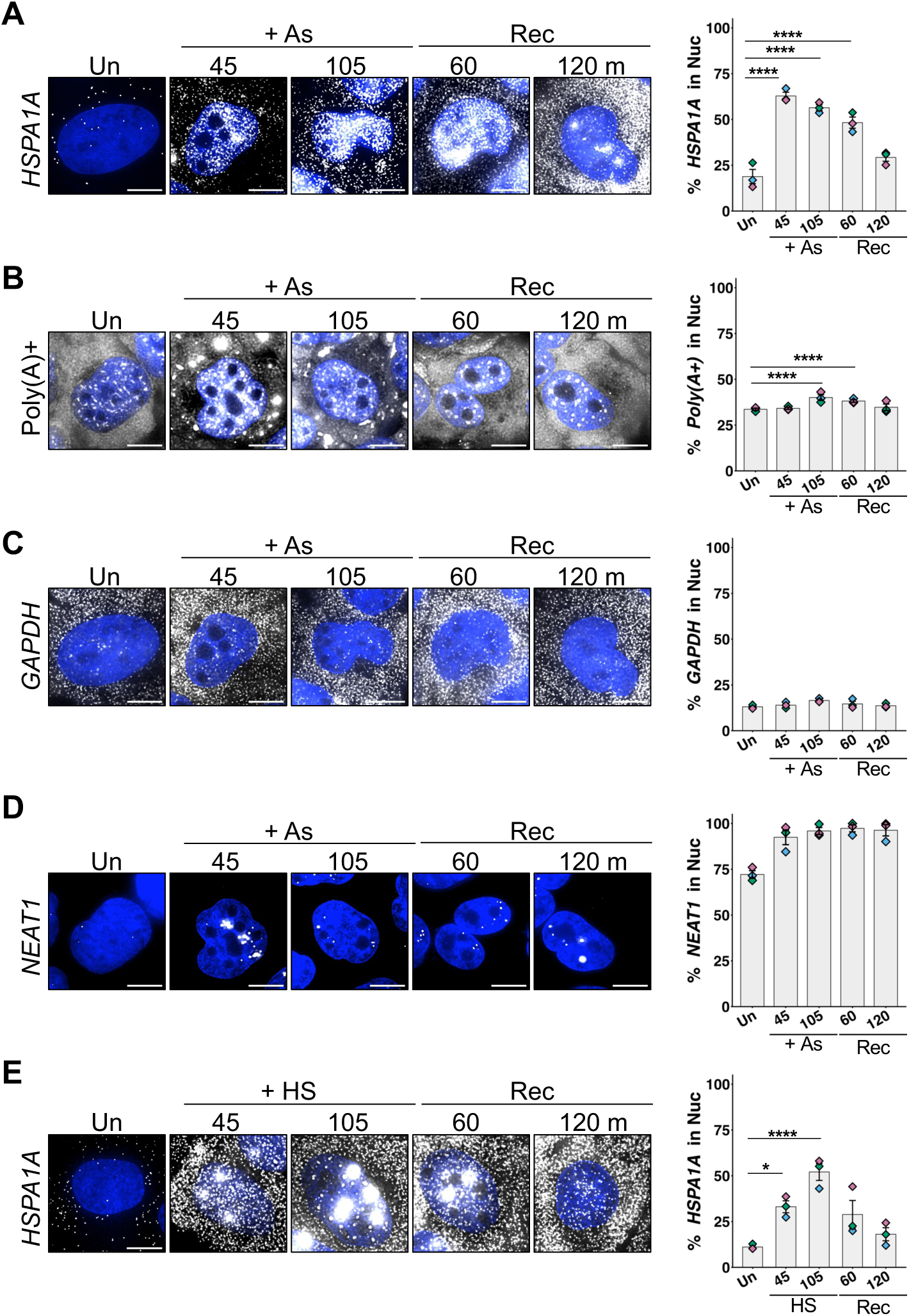
*HSPA1A* accumulates in the nucleus in arsenite and heat stress. Subcellular localization of (A) *HSPA1A*, (B) Poly(A)+ RNAs, (C) *GAPDH*, and (D) *NEAT1* during arsenite stress (“As”, 250 µM; 45 or 105 min), and (E) *HSPA1A* during heat stress (“HS”, 43 °C for 45 or 105 min). *Left:* Representative smFISH images of U-2 OS cells under unstressed (“Un”), stressed, and recovery (“Rec”, 60 or 120 min post-stress) conditions; nuclei in blue and RNA in white. Scale bars: 10 µm*. Right*: Quantification of images from n = 3 independent experiments, with mean +/- s.e.m. of the percent nuclear RNA per cell shown (replicate means shown as diamonds). 46-132 cells were analyzed for each condition. Statistical significance shown using one-way ANOVA tests. (*) *P < 0.05,* (****) *P < 0.001*.

To determine if the nuclear accumulation of *HSPA1A* during stress is generalizable to other stresses that activate the ISR and HSR, we heat stressed cells (at 43°C) under conditions known to upregulate *HSPA1A* (Vera et al. 2014; Lellahi et al. 2018; Theodorakis and Morimoto 1987). We heated cells for 45 or 105 min, or returned them to 37°C (after 45 min heat stress) to recover for 60 or 120 minutes, and performed smFISH for *HSPA1A*. As anticipated, we observed *HSPA1A* was significantly upregulated (∼11-19-fold) during heat stress (**Supplemental Fig. S2E**). We observed that heat stress caused the progressive accumulation of *HSPA1A* in the nucleus after 45 and 105 min of stress (**Figure 1E**). Similarly to arsenite stress, *HSPA1A* was predominantly cytoplasmic after 120 min of recovery from heat stress (**Figure 1E**). We noted that unlike arsenite stress, in which cells exhibited a diffuse localization pattern of *HSPA1A* in the nucleus, *HSPA1A* was diffusely localized and present in large (∼2 µm diameter) and small (∼1 µm diameter) foci in the nucleus in heat-stressed cells (**Figure 1E**). Together, these results demonstrate that the nuclear accumulation of *HSPA1A* occurs across diverse cell types and stresses that induce the HSR.

### *HSPA1A* mRNAs are retained in the nucleus during arsenite and heat stress

High steady-state nuclear *HSPA1A* RNA levels could result from three possible phenomena. First, intense and sustained transcriptional induction that outpaces nuclear export could cause nuclear mRNA accumulation. Second, inhibition of mRNA export could cause nuclear retention of mRNAs. Third, increased cytoplasmic RNA decay rates could result in a higher relative amount of mRNA in the nucleus. To differentiate between these possibilities, we assessed the abundance of *HSPA1A* in the nucleus and cytoplasm over time after shutting off transcription during stress. If *HSPA1A* mRNAs were retained in the nucleus, then the percentage of *HSPA1A* in the nucleus would be unchanged when transcription is shut off. In contrast, if transcripts were not retained in the nucleus, we would expect the percentage of *HSPA1A* in the nucleus to decline, and cytoplasmic mRNA abundance to increase, over time. Alternatively, if *HSPA1A* mRNAs were degraded in the cytoplasm at an increased rate compared to the nucleus, we would expect the number of mRNAs in the cytoplasm to decrease over time.

We stressed cells with arsenite for 45 min, then added the transcription inhibitor actinomycin D or carrier DMSO and fixed cells for smFISH in 15-minute intervals to quantify the percentage of *HSPA1A* in the nucleus and the abundance of *HSPA1A* over time. As expected, actinomycin D effectively repressed *HSPA1A* transcriptional induction during stress (**Supplemental Fig. S3A**). Further, the abundance of *HSPA1A* continually increased from 45 min to 105 min of arsenite stress in cells treated with DMSO, and plateaued after 105 min of stress (**Supplemental Fig. S3B**). We noted that in actinomycin D treated cells, *HSPA1A* was not present in nuclear foci that could represent nuclear speckles and indicate actinomycin D impaired nuclear export, as occurs with longer (4 - 8 hours) transcriptional inhibitor treatments (Williams et al. 2025; Tokunaga et al. 2006). Importantly, there was a ∼50% reduction in the percentage of *HSPA1A* mRNAs in the nucleus between 0 and 30 min of actinomycin D treatment. These results suggest that *HSPA1A* is released from the nucleus up to 75 min post-arsenite stress (**Figure 2A&B**). The percentage of *HSPA1A* in the nucleus was sustained at ∼25% between 30 and 60 min of actinomycin D treatment, suggesting that *HSPA1A* is retained in the nucleus after 75 min of arsenite stress (**Figure 2A&B**). Together, these results suggest that nuclear *HSPA1A* export is progressively inhibited during arsenite stress.

**Figure 2.**
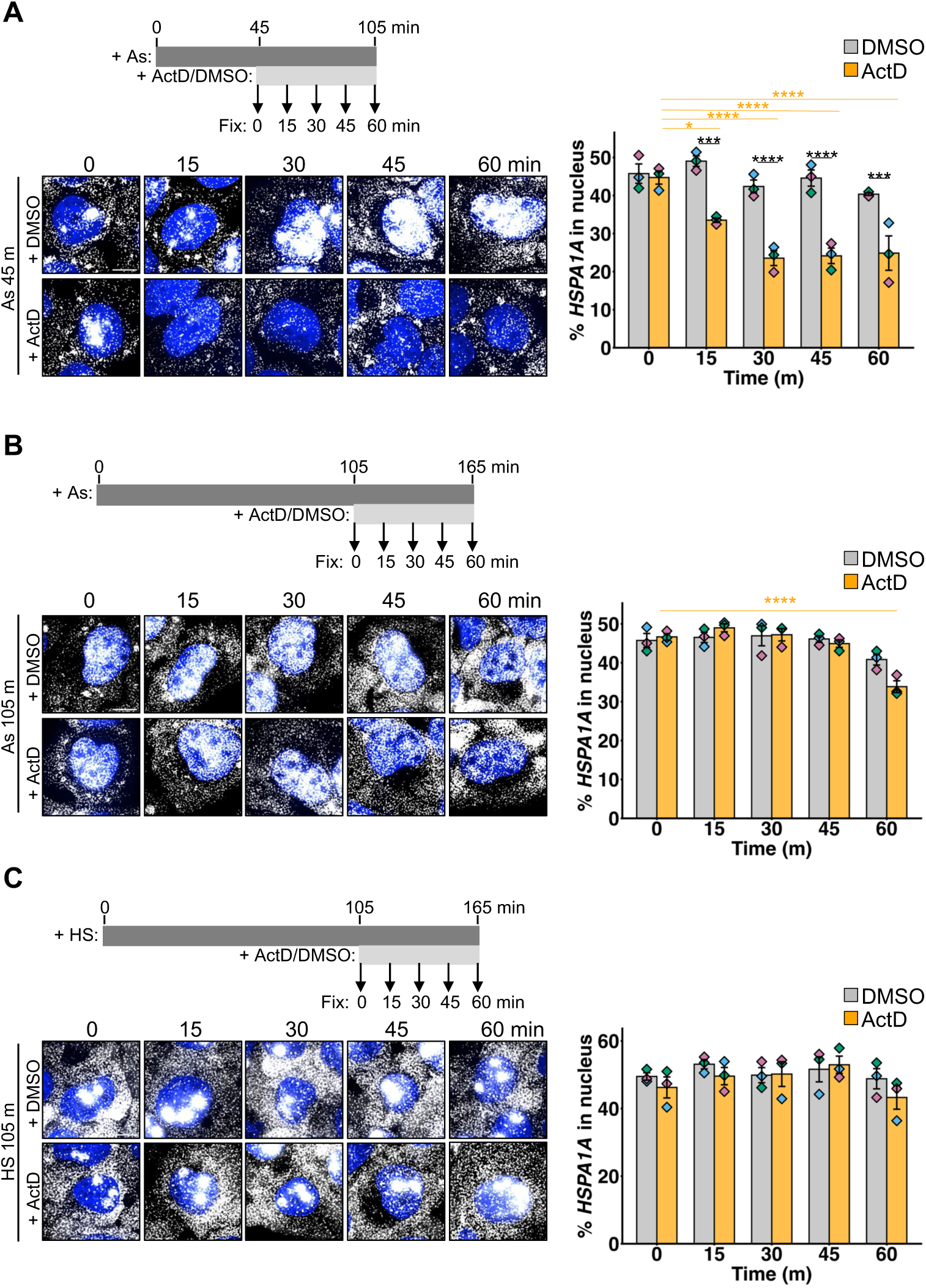
*HSPA1A* nuclear export is progressively inhibited during arsenite and heat stress. (A) The subcellular localization of *HSPA1A* mRNAs from smFISH images collected every 15 min in cells treated with actinomycin D (“ActD”) or carrier control DMSO (0.1%) starting at 45 min of arsenite stress (“As”, 250 µM). *Left, top:* Experimental timeline. *Left, bottom:* Representative smFISH images; nuclei in blue and *HSPA1A* in white. *Right*: Quantification of images with the mean +/- s.e.m. of the percentage of *HSPA1A* in the nucleus shown; grey bars indicate DMSO treatment, and orange bars indicate ActD treatment. Diamonds represent the mean of each independent replicate (n = 3). (B) As in (A), with ActD added at 105 min arsenite stress. (C) As in (B), with cells heat stressed (“HS”; 43 °C) for 105 min. prior to ActD addition. One-way ANOVAs were performed to assess statistical significance, with orange lines representing significance between ActD conditions and black indicating significance between DMSO and ActD treatments. (*) *P < 0.05,* (***) *P < 0.005,* (****) *P < 0.001.* Scale bars: 10 µm; 69-98 cells counted per condition.

The progressive nuclear retention of *HSPA1A* during early arsenite stress suggests a time-dependent inhibition of nuclear mRNA export during stress. We therefore hypothesized *HSPA1A* export from the nucleus would be inhibited following longer stress treatments. To test this hypothesis, we stressed cells with arsenite for 105 min and added actinomycin D or DMSO, and fixed cells in 15-minute intervals to measure the percentage of *HSPA1A* in the nucleus and total *HSPA1A* abundance. We again confirmed actinomycin D shut off transcription by measuring *HSPA1A* levels in a parallel treatment (**Supplemental Fig. S3C**). A key observation is that the percentage of *HSPA1A* in the nucleus was largely unchanged in cells treated with actinomycin D beginning at 105 min of arsenite stress (**Figure 2B**). We observed a small ∼13 percentage point decline in *HSPA1A* in the nucleus at the final time point (165 min post-stress) in cells treated with actinomycin D, suggesting a limited (yet severely reduced) amount of export occurs at this late stress stage. The levels of *HSPA1A* did not change in the DMSO-treated control over time, suggesting that *HSPA1A* transcription had ceased with export inhibition by 105 min of arsenite stress (**Supplemental Fig. S3D**). Together, these observations indicate that arsenite stress causes a reversible, time-dependent inhibition of *HSPA1A* nuclear export.

To evaluate whether *HSPA1A* retention in the nucleus is a shared feature of other stresses, we next tested whether *HSPA1A* accumulates in the nucleus when transcription is inhibited during heat stress. We stressed cells at 43°C for 105 min, added either actinomycin D or DMSO, and fixed cells for smFISH in 15-minute intervals for 60 min. In parallel, we confirmed actinomycin D inhibited the transcription of *HSPA1A* during heat stress (**Supplemental Fig. S3E**). Similar to prolonged arsenite stress, we found that the percentage of *HSPA1A* in the nucleus did not change over the actinomycin D time course (**Figure 2C**). We observed 46% +/- 3.1% *HSPA1A* in the nucleus at 105 min, and 43% +/- 3.4% was nuclear at 165 min of heat stress (**Figure 2C**). Also similar to arsenite stress, the levels of *HSPA1A* did not change after 105 min of heat stress (**Supplemental Fig. S3F**). These data demonstrate that *HSPA1A* accumulation in the nucleus during arsenite and heat stress results from the retention of these RNAs in the nucleus, indicative of impaired nuclear mRNA export.

### Export of stress-induced gene mRNAs is inhibited during arsenite stress

We next addressed whether nuclear retention is specific to *HSPA1A*, other transcriptionally-induced RNAs, or all RNAs during stress. We therefore identified the nuclear-enriched transcriptome during arsenite stress. We stressed cells with arsenite for 105 min, then collected RNA from whole cell lysates or from nuclear and cytoplasmic fractions. We then performed RNA sequencing of rRNA-depleted samples. We verified successful cell fractionation by demonstrating the nuclear lncRNA *MALAT1* was enriched in nuclear fractions and depleted in cytoplasmic fractions, and the constitutively expressed *GAPDH* mRNA was depleted in nuclear fractions and enriched in cytoplasmic fractions using RT-qPCR (**Supplemental Fig. S4A**). Further, we confirmed there was a higher proportion of intronic RNA-seq reads in nuclear lysates than cytoplasmic lysates (**Supplemental Fig. S4B**). As additional validation, we compared the nuclear and cytoplasmic reads to assess the degree of partitioning for each transcript in unstressed cells, which showed nuclear lncRNAs *MALAT1* and *NEAT1* were highly enriched in the nucleus while *GAPDH* and other constitutively expressed mRNAs were highly cytoplasmic, as expected (**Supplemental Fig. S4C**).

We next identified the transcripts that were induced by arsenite stress to begin to address whether these transcripts were specifically retained in the nucleus. We determined the transcriptome-wide changes in RNA abundance between stressed and unstressed samples, and identified 1,561 significantly upregulated protein-coding mRNAs (≥1.5 fold, p<0.05; **Figure 3A**). Significantly upregulated protein-coding transcripts included *PNLDC1*, *HSPA1A*, *GADD34*, *HMOX1* and *FOS* (**Figure 3A**). Upregulation of *FOS* and other immediate-early genes (e.g., *JUN*, *EGR2*) was likely due to the arsenite stress and not serum in the media, as media was also exchanged on unstressed cells prior to collection. We observed fewer downregulated transcripts, as 1,115 RNAs protein coding transcripts were significantly reduced in abundance (≥1.5-fold) upon arsenite stress (**Figure 3A**). The levels of *GAPDH* and *NEAT1* did not change upon stress (**Figure 3A**). Therefore, we defined the pool of mRNAs that are significantly upregulated during prolonged arsenite stress as stress-induced transcripts.

**Figure 3.**
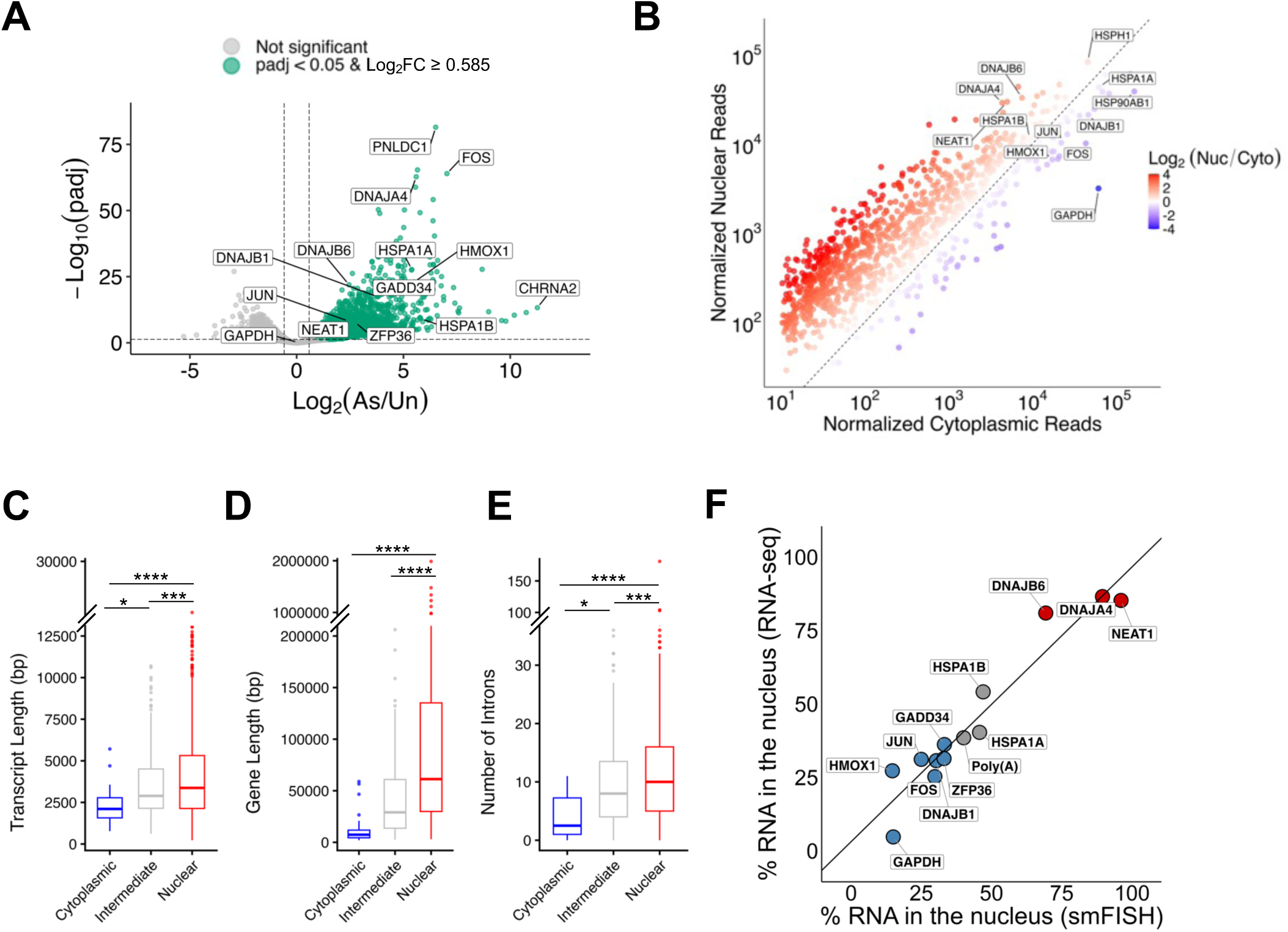
Transcriptome-wide analysis of nuclear and cytoplasmic stress-induced mRNAs during arsenite stress. (A) Volcano plot showing differential gene expression between unstressed (“Un”) and arsenite stressed (“As”; 250 µM; 105 min) cells. Green dots indicate significantly upregulated genes (padj < 0.05 & Log2FC ≥ 0.585); n = 3 independent replicates. (B) Scatter plot of normalized read counts showing nuclear and cytoplasmic distribution of upregulated, protein-coding RNAs during arsenite stress (250 µM; 105 min) with labels indicating representative stress-induced transcripts and *GAPDH* and *NEAT1* as cytoplasmic and nuclear fiduciaries. Color-coded for Log2(Nuc/Cyto) ratio, with red representing nuclear-enrichment and purple representing cytoplasmic enrichment; n=3 independent replicates. (C-E) Boxplots (median, 25th and 75th percentiles shown) of transcript length (C), gene length (D), and intron count (E) for cytoplasmic-enriched (Log2(Nuc/Cyto) ≤ -0.585), nuclear-enriched (Log2(Nuc/Cyto) ≥ 0.585), and intermediate genes as defined in (B). Wilcoxon rank sum test was used to assess statistical significance. (****) *P < 0.001.* (F) Correlation between the percentage of each representative RNA in the nucleus determined using RNA-seq versus smFISH.

We next evaluated whether stress-induced transcripts preferentially accumulated in the nucleus by comparing nuclear to cytoplasmic read counts and assessing if genes that were upregulated during stress were distributed unevenly across compartments. Notably, we observed a strong enrichment in stress-induced genes in the nuclear compartment (**Figure 3B**). Stress-induced mRNAs were more likely to be enriched in the nucleus (by ∼4.2-fold on average) and constitutively expressed transcripts were more likely to be enriched in the cytoplasm (by ∼1.5-fold on average). We observed 894 stress-induced transcripts were significantly enriched in the nucleus and also significantly upregulated by ≥1.5-fold (**Figure 3B; Supplementary Table S1**). These nuclear mRNAs included HSP40-encoding transcripts *DNAJA4* and *DNAJB6*, and enrichment analysis revealed HSF1 binding motifs were over-represented (p-adj = 5.24 x 10^-8^). As expected, and similar to unstressed cells (**Supplemental Fig. S4C**), the nuclear lncRNA *NEAT1* was enriched in the nuclear fraction and constitutively expressed mRNA *GAPDH* was enriched in the cytoplasm (**Figure 3B**). Thus, the majority of stress-induced transcripts are enriched in the nucleus during arsenite stress.

We next asked if any stress-induced mRNAs had accumulated in the cytoplasm during arsenite stress, which could suggest specificity in nuclear mRNA retention. We observed 55 stress-induced gene mRNAs that were enriched in the cytoplasm, which included *HMOX1*, *FOS*, *JUN*, and HSP40 and HSP90 mRNAs (**Figure 3B; Supplementary Table S1**). Stress-induced transcripts enriched in the cytoplasm were shorter than nuclear-enriched transcripts (**Figure 3C**), and were encoded by shorter genes (**Figure 3D**) that had fewer introns (**Figure 3E**). We verified that the enrichment of mRNAs in nuclear or cytoplasmic compartments tightly correlated with smFISH results (R^2^=0.917), demonstrating that nucleocytoplasmic fractionation RNA-seq provides a complementary approach to measure global changes in RNA localization (**Figure 3F**). These results indicate that most stress-induced mRNAs are enriched in the nucleus and a subset of these mRNAs escapes the nucleus during arsenite stress.

### Nuclear mRNA export is progressively inhibited during stress

The progressive decline in the nuclear export of *HSPA1A* during stress, and the observation that most stress-induced mRNAs are enriched in the nucleus, suggested a temporal gating model in which mRNAs that are transcriptionally induced early during stress would escape into the cytoplasm before nuclear export was inhibited. An alternative explanation is that some stress-induced gene mRNAs are preferentially exported from the nucleus through their interactions with export licensing factors, as would be expected based on the current paradigm of stress-induced mRNA export (Seidler and Sträßer 2024).

To differentiate between these two possibilities, we first monitored the nuclear export of stress-induced mRNAs. We used actinomycin D to shut off transcription at 105 min of arsenite stress and assessed the nuclear:cytoplasmic mRNA ratio of three types of transcripts: those that are cytoplasm-enriched (*FOS*, *HMOX1*, *DNAJB1*, *JUN*, and *ZFP36*), equally distributed (*GADD34* and *HSPA1B)*, or nuclear-enriched (*DNAJA4* and *DNAJB6*). We also evaluated total poly(A)+ RNA using oligo(dT) FISH. We verified that the actinomycin D successfully shut off transcription, and in the process, confirmed all candidate mRNAs were transcriptionally induced during arsenite stress (**Supplemental Fig. S5**). If cytoplasm-enriched mRNAs have some property that allows them to escape nuclear retention, we anticipated that the percentage of these mRNAs in the nucleus would decline over time after the addition of actinomycin D.

Importantly, we observed the nuclear export of every tested mRNA was inhibited during late arsenite stress (**Figure 4**). The nuclear export of cytoplasm-enriched mRNAs was inhibited during late stress (**Figure 4A**). We confirmed that *FOS* and *ZFP36* underwent more efficient export from the nucleus during early (45 min) stress than late (105 min.) stress (**Supplemental Fig. S6**; **Figure 4A**). Similarly, the percentage of poly(A)+ RNA in the nucleus was unchanged in cells treated with actinomycin D during late arsenite stress (**Figure 4B**). Furthermore, the percentage of nuclear transcripts that were equivalently distributed in the nucleus and cytoplasm (*HSPA1B* and *GADD34*; **Figure 4C**) or enriched in the nucleus (*DNAJA4* and *DNAJB6*) (**Figure 4D**) did not change during the actinomycin D time course. Transcription of all candidate stress-induced genes ceased by 105 min, as there were no significant increases in any of these mRNAs in the DMSO-treated cells at up to 165 minutes of arsenite stress (**Supplemental Fig. S5**). We ruled out the possibility that differential cytoplasmic RNA decay contributes to the nuclear accumulation of these transcripts, as the total abundance of the transcripts did not decrease over time during actinomycin D treatment (**Supplemental Fig. S5**). Altogether, these results indicate that stress-induced mRNAs are generally retained in the nucleus during stress, irrespective of transcript-specific licensing factors.

**Figure 4.**
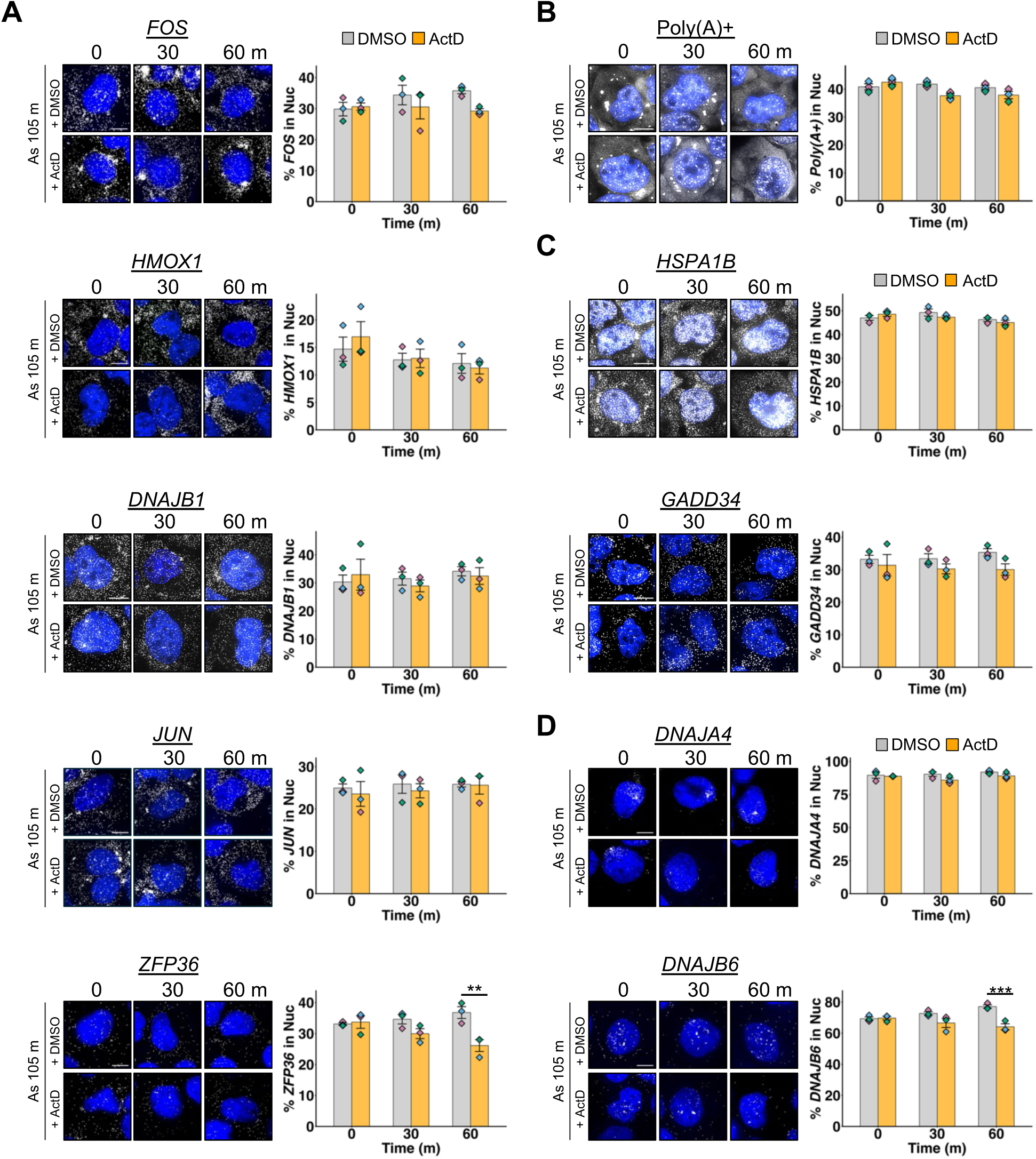
Stress-induced mRNAs are retained in the nucleus during arsenite stress. Actinomycin D treatments (5 µg/mL) compared to DMSO carrier controls (0.1%) were done starting at 105 min arsenite stress (250 µM) and smFISH performed to detect (A) cytoplasm-enriched stress-induced mRNAs, (B) poly(A)+ RNA, (C) intermediate mRNAs, and (D) nuclear-enriched mRNAs. Samples were collected in 30-min intervals. *Left:* Representative images with nuclei in blue and RNAs in white; scale bars 10 µm. *Right:* Quantification of the mean +/- s.e.m. nuclear percentage over time for each RNA, with grey representing DMSO treatment and orange representing ActD treatment. Diamonds indicate the average of each independent (n = 3) replicate. (B) One-way ANOVAs were performed to assess statistical significance. (**) *P < 0.01,* (***) *P < 0.005.* 51-119 cells counted per condition.

### Temporal gating underlies nucleocytoplasmic mRNA distribution during stress

Our results point to a temporal gating model wherein mRNAs that are transcribed early during stress are efficiently exported into the cytoplasm before nuclear mRNA export is suppressed. To begin to interrogate this model, we first determined if mRNAs transcribed early during stress were more likely to be enriched in the cytoplasm. To identify transcripts that were transcribed early or late during stress, we performed poly(A)-enriched RNA sequencing from samples collected over a time course of arsenite stress (0, 15, 30, 45, 75, and 105 min) (**Figure 5A**). We then determined which mRNAs increased in abundance at each time point (≥ 50% increase and p < 0.05), then binned them as early- (genes upregulated between 0 and 45 min), or late- (genes upregulated between 45 and 105 min) stress-induced genes. We identified 738 “early” genes, including *ATF3*, *FOS*, *JUN*, *HMOX1*, and *DNAJB1* (**Figure 5B**). Additionally, we found 652 "late” genes, including *DNAJA4*, *HSPH1*, and *BAG3* that increased in abundance starting at 75 min of stress (**Figure 5C; Supplemental Fig. S7**). Genes upregulated in both phases (e.g., *HSPA1A* and *HSPA1B)* were classified as “early”, although many exhibited sustained induction over time and appeared upregulated in both early- and late- stress datasets.

**Figure 5.**
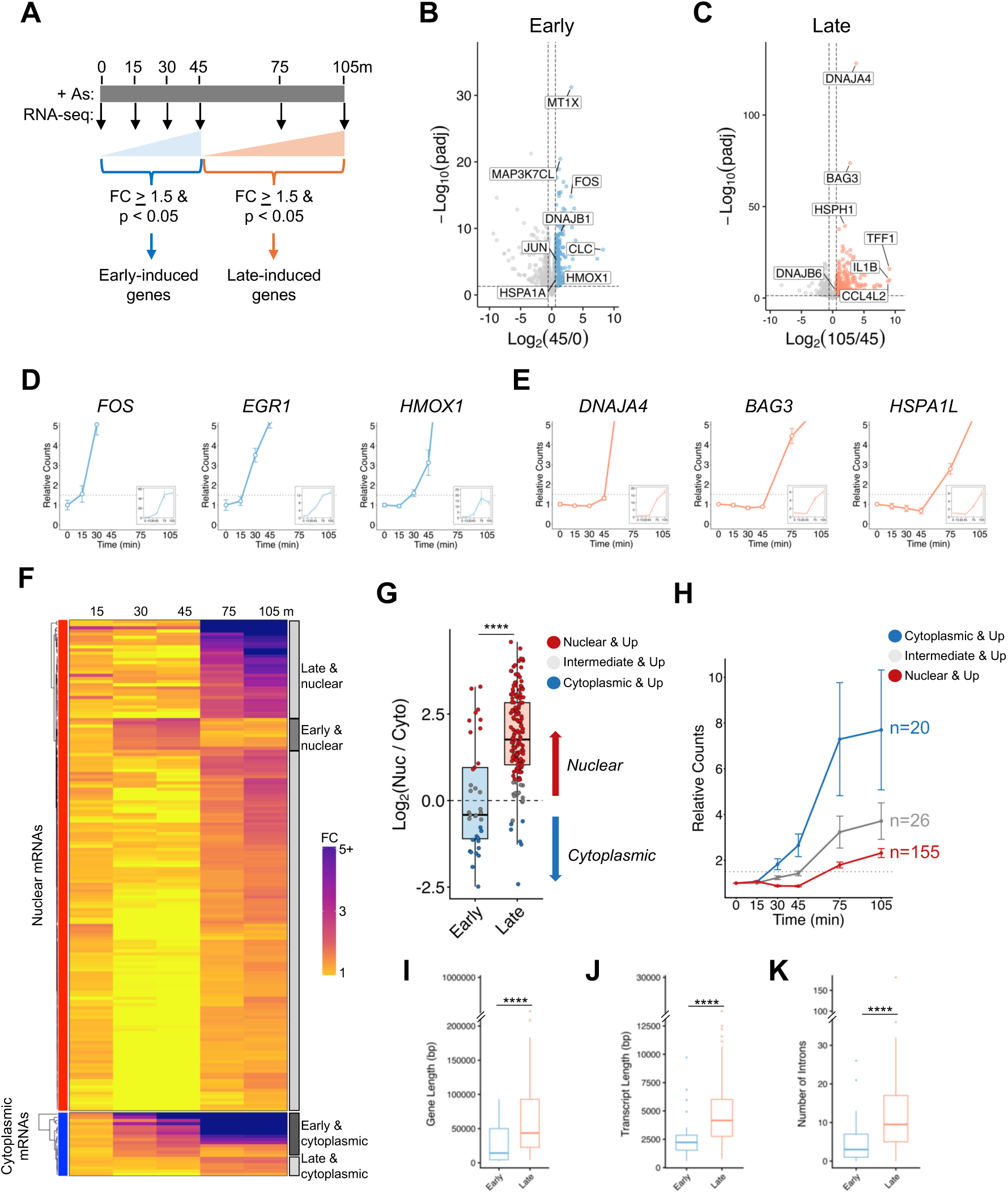
Early-transcribed mRNAs are exported from the nucleus while late-transcribed mRNAs are retained in the nucleus during arsenite stress. (A) Experimental timeline and filters to identify early- and late- stress-induced genes during arsenite (“As”, 250 µM) stress; n=3 independent replicates. (B) Differential gene expression between unstressed (0 min) and early (45 min) arsenite stress. Blue dots represent significantly upregulated genes (padj < 0.05; Log2FC ≥ 0.585). (C) Differential gene expression between 45 and 105 min of arsenite stress. Orange dots represent significantly upregulated genes (padj < 0.05; Log2FC ≥ 0.585). (D) Normalized counts relative to unstressed (0 min) for representative early-induced transcripts. Y-axes are shrunk to visualize when counts cross the 1.5 FC threshold; insets show full-scale y-axes. Average normalized counts +/- s.e.m. for each time point shown. (E) As in (D), but for representative late-induced transcripts. (F) Heatmap of mRNAs represented in nuclear/cytoplasmic fractionation datasets (Figure 3) and the early-late RNA-sequencing datasets. Data grouped by compartment (red for nuclear genes and blue for cytoplasmic) and hierarchical clustering. *Right*: Categorization of genes into early, late, cytoplasmic, and nuclear. (G) Boxplots (median, 25th and 75th percentiles) of the Log2(Nuc/Cyto) for early and late genes. Red dots represent nuclear-enriched genes (Log2(Nuc/Cyto) ≥ 0.585), blue dots represent cytoplasm-enriched genes (Log2(Nuc/Cyto) ≤ -0.585), and grey dots represent intermediate genes. (H) Average +/- s.e.m. of normalized counts over time relative to unstressed samples for nuclear-enriched, cytoplasm-enriched, and intermediate gene classes. (I-K) Boxplots (median, 25th and 75th percentiles) of gene length (I), transcript length (J), and intron count (K) for early-induced and late-induced genes. Wilcoxon rank sum test done to assess statistical significance. (****) *P < 0.001*.

We next compared the early- and late- induced transcripts to their nucleocytoplasmic partitioning. If the temporal gating model were supported, then mRNAs upregulated early would be more likely to be enriched in the cytoplasm (e.g. *FOS*) than mRNAs upregulated at later time points, while late-induced mRNAs would be enriched in the nucleus. We categorized the stress-induced transcripts as either “nuclear” or “cytoplasmic” if they were ≥1.5-fold increased in either compartment, and performed four types of analyses. Analyses were restricted to transcripts that were significantly upregulated between 0 and 105 min of stress and present in the nuclear:cytoplasmic fractionation dataset. This resulted in a subset of 201 genes, of which 37 were classified as “early” and 164 were classified as “late”. For example, immediate-early genes *FOS*, *EGR1*, and *HMOX1* were upregulated early (**Figure 5D**) and were localized to the cytoplasm during stress. In contrast, HSPs *DNAJA4* and *HSPA1L*, and co-chaperone *BAG3* were upregulated late (**Figure 5E**), and were localized to the nucleus.

In our first analysis, we visualized the abundance of each nuclear- or cytoplasm- enriched stress-induced transcript at each time point relative to the unstressed (“0 min”) condition as a heat map. Importantly, the majority of nuclear-enriched mRNAs were upregulated on or after 75 min post-arsenite stress (“late”), and the cytoplasm-enriched mRNAs were mostly upregulated within 45 min (“early”) (**Figure 5F**). Some exceptions were present. Eleven “early” mRNAs were nuclear-enriched. These mRNAs had lower read depth, were encoded by longer genes with more introns, and had higher relative intronic reads than other early-induced mRNAs, consistent with nuclear retention via intron detention (**Supplemental Fig. S6A**; **S8**). Six “late” mRNAs were cytoplasm-enriched, but were already highly cytoplasmic before stress (**Supplemental Fig. S4C**) and did not increase in cytoplasmic abundance during stress, suggesting their nascent transcripts are nuclear-retained. One of these mRNAs, *ANKRD1*, exhibited stress-dependent intronic read accumulation that may also indicate an intron detention mechanism of nuclear retention (**Supplemental Fig. S9**). Overall, the timing of stress-induced gene upregulation was strongly associated with nucleocytoplasmic partitioning, with early induction corresponding to cytoplasmic enrichment and late induction corresponding to nuclear enrichment.

Second, we asked whether early-induced mRNAs as a class were more cytoplasmic than late-induced mRNAs. We found that early-induced transcripts were more cytoplasmic (median 1.33-fold cytoplasmic enrichment) than late-induced transcripts (median 3.43-fold nuclear enrichment) (**Figure 5G**). This analysis included mRNAs that were not significantly enriched in the nucleus or cytoplasm, but were upregulated early or late during stress. The same few transcripts that were exceptions to these associations were classified as “early nuclear” or “late cytoplasmic” in **Figure 5F** as described above. Thus, mRNAs transcribed early upon stress were more likely to accumulate in the cytoplasm than late-transcribed mRNAs.

Third, we performed a trajectory analysis of all stress-induced genes by assessing the relative abundance of cytoplasmic, nuclear, and intermediate localized mRNAs in aggregate over the stress time course. This analysis revealed that cytoplasmic-enriched mRNAs were upregulated by ∼1.83-fold within just 30 min of stress. In contrast, nuclear-enriched mRNAs required 75 min of stress to reach a similar ∼1.8-fold increase. Intermediate genes mirrored their balanced localization with an ‘intermediate’ induction profile, increasing from 1.2 fold at 30 minutes and 3.17-fold at 75 minutes (**Figure 5H**). Furthermore, following 105 min of stress, cytoplasmic genes exhibited the most robust induction (7.7-fold), compared to intermediate genes (3.7-fold), and nuclear-enriched genes (2.3-fold) (**Figure 5H**). Therefore, when examined in aggregate, the temporal profiles of transcriptional induction reflected nucleocytoplasmic partitioning during stress.

Fourth, we asked if the early mRNAs were shorter, were encoded by shorter genes, and had fewer introns, which would enable more rapid transcription and processing than longer, intron-dense mRNAs (estimated as ∼1 - 4 kb/min (Singh and Padgett 2009; Maiuri et al. 2011)). Compared to late-induced genes, we found that early-induced mRNAs were encoded by shorter genes (**Figure 5I**), their transcripts were shorter (**Figure 5J**) and these genes contained fewer introns (**Figure 5K**). Therefore, the majority of stress-induced transcripts that are exported into the cytoplasm are expressed early during stress, while those enriched in the nucleus are transcribed later. Together, these data support a temporal gating model where early transcriptional induction facilitates mRNA release from the nucleus before the stress-induced blockade of nuclear export.

### Nuclear retention of an inducible reporter RNA supports the temporal gating model

We next directly tested the temporal gating model by monitoring the nuclear export kinetics of doxycycline (dox)- induced reporter mRNAs. If the temporal gating model were supported, we expected the reporter RNA would progressively increase in the nucleus and nuclear export would decline over time during arsenite and heat stress. We used a previously generated U-2 OS cell line in which the *GADD34* 5’ untranslated region is upstream of *Renilla* luciferase (*GADD34-Luc*), and detected the mRNAs using smFISH probes to the luciferase mRNA (Helton et al. 2025). We first established the time it takes for the onset of transcription and for the cytoplasmic accumulation of *GADD34-Luc* reporter mRNAs. We added dox at 15-minute intervals and monitored the number of cells with smFISH spots and the abundance of *GADD34-Luc* spots in the cytoplasm. We established that it takes 30 min for the onset of *GADD34-Luc* transcriptional induction (**Figure 6A; Supplemental Figure S10A**). Importantly, the number of *GADD34-Luc* mRNAs localized to the cytoplasm significantly increased starting at 45 min post-dox addition, with a ∼5-fold increase observed at 60 min when ∼50% of the *GADD34-Luc* mRNAs were cytoplasmic (**Figure 6B**; **Supplemental Fig. S10A**). These results indicate that the *GADD34-Luc* reporter mRNA is transcribed after a 30 min lag time and accumulates in the cytoplasm beginning 45 min after dox addition (**Figure 6C**), demonstrating we can rapidly induce transcription and monitor reporter mRNA accumulation in the cytoplasm.

**Figure 6.**
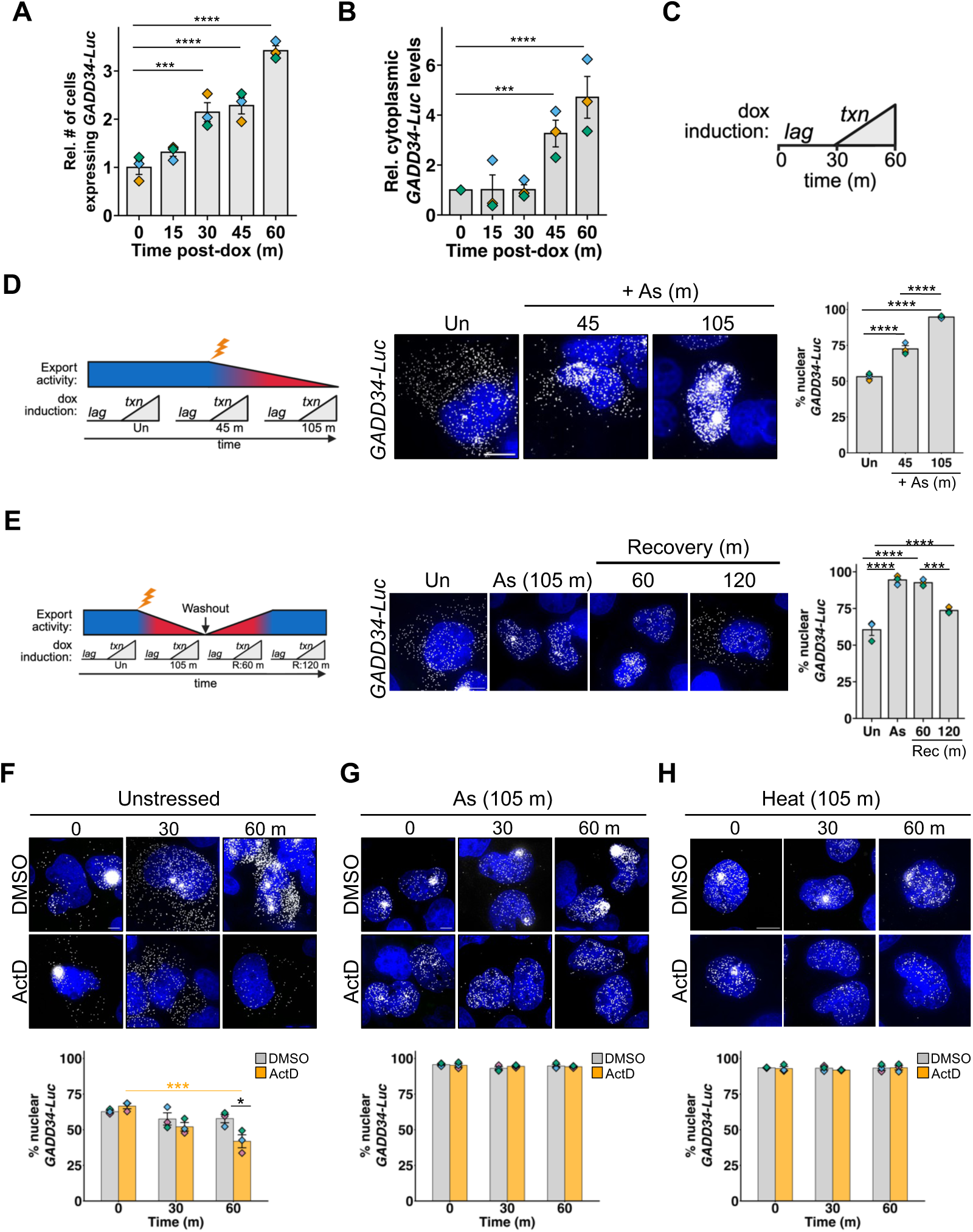
Nuclear export of inducible *GADD34-Luc* reporter mRNAs is progressively inhibited during arsenite and heat stress. (A) The relative number of cells with *GADD34-Luc* smFISH spots up to 60 min after doxycycline (“dox”; 1 µg/mL) treatment (215-524 cells per image). (B) Relative cytoplasmic *GADD34-Luc* smFISH spots per cell up to 60 min after dox addition. (C) Schematic of *GADD34-Luc* transcription (“txn”) 30 min after dox addition. (D) *Left:* Experimental paradigm in relation to export activity over time during stress. Cells were unstressed or stressed with arsenite (“As”; 250 µM; lightning bolt) for 45 or 105 min. and smFISH to *GADD34-Luc* performed. *Middle*: Representative images of *GADD34-Luc* (white) and nuclei (blue). *Right:* Quantification of the percentage of *GADD34-Luc* in the nucleus (63-83 cells counted per condition). (E) Subcellular localization of *GADD34-Luc* during and after recovery from arsenite stress. *Left:* Experimental schematic in relation to nuclear mRNA export activity before, during, and after washout of arsenite stress (105 min; “As”; 250 µM; lightning bolt). *Middle:* Representative images of *GADD34*-*Luc* mRNAs (white) and nuclei (blue). *Right:* Quantification of nuclear *GADD34-Luc* percentage in each condition (50-54 cells counted per condition). (F) smFISH of *GADD34-Luc* in unstressed cells at 0, 30, or 60 min post-actinomycin D (“ActD”; orange bars) addition compared to carrier control DMSO (0.1%; grey bars). *Top*: Representative images of *GADD34-Luc* (white) and nuclei (blue). *Bottom:* Quantification of the nuclear *GADD34-Luc* percentage over time (69-99 cells counted per condition). (G) As in F, except cells were treated with arsenite for 105 min prior to ActD or DMSO addition (48-58 cells counted per condition) (H) As in H, except cells were heat stressed (“HS”; 43°C for 105 min) before DMSO or ActD treatment (53-75 cells counted per condition). Cells were treated with dox for 60 min before ActD or DMSO treatment for F-H. Scale bars in images: 10 µm. Quantifications represent average +/- s.e.m. from n = 3 independent replicates, with individual replicate means shown as diamonds. One-way ANOVAs were done to determine statistical significance across time points, with orange indicating significance between ActD time points and black indicating significance between DMSO and ActD time points. (*) *P < 0.05,* (***) *P < 0.005,* (****) *P < 0.001*.

We next assessed whether the *GADD34-Luc* reporter mRNA accumulated in the nucleus during arsenite stress. We treated cells with dox and either left them unstressed, or stressed them with arsenite for 45 or 105 min (**Figure 6D**). In all cases, the cells were treated with dox for 60 min prior to fixation. Consistent with the temporal gating model, we observed that the reporter mRNAs accumulated in the nucleus over time during stress (**Figure 6D**; **Supplemental Fig. S10B**). While unstressed cells had 53% +/- 1.4% mRNA in the nucleus, likely owing to robust dox-induced transcription, 73% +/- 2.3% of the *GADD34-Luc* mRNA was in the nucleus in cells stressed for 45 min (**Figure 6D**). Although the dox was added 15 minutes prior to arsenite addition, it is unlikely that the ∼30% of *GADD34-Luc* present in the cytoplasm was exported before stress was added because very few mRNAs were observed in the cytoplasm at 30 min post-dox addition (**Figure 6B&C**). Cells stressed for 105 min exhibited 95% +/- 0.5% *GADD34-Luc* mRNA in the nucleus (**Figure 6D**). Thus, inducible reporter mRNAs progressively accumulate in the nucleus during arsenite stress.

Importantly, we found evidence that the nuclear export of *GADD34-Luc* mRNAs was restored after stress. Cells treated with arsenite for 105 min displayed 95% +/-1.7% nuclear *GADD34-Luc*, while cells recovering from stress for 120 min after arsenite washout exhibited 74% +/- 1.2% nuclear *GADD34-Luc*, similar to unstressed cells (**Figure 6E**). The observations that *GADD34-Luc* was not downregulated during arsenite stress, and that cytoplasmic *GADD34-Luc* levels increased during recovery from stress ruled out the possibility that arsenite inhibited *GADD34-Luc* transcription (**Supplemental Fig. 10C&D**). The observed progressive accumulation of *GADD34-Luc* in the nucleus during arsenite stress, and cytoplasmic accumulation after stress removal, support the hypothesis that nuclear export is generalizable and reversibly inhibited during arsenite stress.

We next tested whether *GADD34-Luc* mRNAs are retained in the nucleus during heat or arsenite stress by controlling for transcriptional induction and accounting for changes in mRNA levels due to decay. We performed actinomycin D shutoff experiments and assessed the percentage of *GADD34-Luc* in the nucleus in cells that were unstressed or stressed with heat or arsenite for 105 min (**Figure 6F-H**). In all conditions, cells were treated with dox for 60 min before addition of actinomycin D. We first assessed *GADD34-Luc* export in unstressed cells. The percentage of *GADD34-Luc* in the nucleus in unstressed cells declined significantly over time, demonstrating the reporter was exported to the cytoplasm in unstressed cells and indicating that actinomycin D did not inhibit nuclear mRNA export (**Figure 6F**). Further, we observed a reduction of *GADD34-Luc* levels in the cytoplasm and in the entire cell over time, and the calculated mRNA half-life of 34 +/- 3.1 min indicates this transcript is rapidly degraded under basal conditions (**Supplemental Fig. S10D&E**). Given that this mRNA is unstable and mRNA decay primarily occurs in the cytoplasm, the observed ∼40% reduction in nuclear RNA levels by 60 min post-actinomycin D substantially underestimates the actual mRNA export rate in unstressed cells. Therefore, *GADD34-Luc* is efficiently exported from the cytoplasm in unstressed conditions.

In contrast to unstressed cells, *GADD34-Luc* mRNA levels persisted in the nucleus in arsenite-stressed cells treated with actinomycin D (**Figure 6G**). The percentage of *GADD34-Luc* in the nucleus in arsenite-stressed cells was 95.5% +/- 1.3% at time 0, and remained high at 94.4% +/- 0.98% 60 min after actinomycin D addition. No cytoplasmic decay was observed for arsenite-stressed cells, as the number of mRNAs per cell did not significantly change after actinomycin D treatment (**Supplemental Fig. S10F**). This was not specific to arsenite stress, as *GADD34-Luc* also persisted in the nucleus after 105 min of heat stress (**Figure 6H**). Specifically, *GADD34-Luc* was 93% +/- 1.4% at time 0, and 94% +/- 1.5% after 60 min of actinomycin D treatment during heat stress. Similarly to arsenite stress, the levels of *GADD34-Luc* did not change during the heat stress time course after actinomycin D treatment (**Supplemental Fig. S10G**). These findings demonstrate that newly transcribed mRNAs are retained in the nucleus during arsenite and heat stress regardless of their promoter or whether they are exogenous or endogenous genes. Taken together, the results of these reporter assays support the temporal gating model and demonstrate early transcribed mRNAs are exported while later transcribed mRNAs are retained in the nucleus.

## Discussion

The selective nuclear export of stress-induced transcripts has long been ascribed to transcript-specific elements that license them for export while other mRNAs are retained in the nucleus (Saavedra et al. 1996, 1997; Coban et al. 2024; Zander et al. 2016). Our findings challenge this model and instead support a temporal gating mechanism wherein the timing of transcription dictates whether or not stress-induced transcripts are exported to the cytoplasm during arsenite or heat stress. Transcriptome-wide assays revealed that most stress-induced gene mRNAs are retained in the nucleus during arsenite stress, while a subset of stress-induced transcripts are enriched in the cytoplasm. Importantly, while all tested mRNAs could export from the nucleus early during stress, no mRNAs were efficiently exported during late stress. Global gene expression profiling over time demonstrated that the earliest transcribed mRNAs accumulate in the cytoplasm, while late-transcribed mRNAs persist in the nucleus during stress. Nucleocytoplasmic partitioning of inducible reporter mRNAs demonstrated the progressive nuclear accumulation of newly transcribed RNAs as stress proceeded. Evaluation of the nuclear mRNA export kinetics of the inducible reporter transcripts directly demonstrated the loss of nuclear export capacity during arsenite and heat stress. Together, these observations indicate that mRNAs that are transcribed early in the stress response escape the nucleus, while later transcribed mRNAs are not transported into the cytoplasm. We therefore propose a temporal gating model that represents the progressive inhibition of nuclear mRNA export during stress and defines the cytoplasmic stress-induced transcriptome (**Figure 7**).

**Figure 7.**
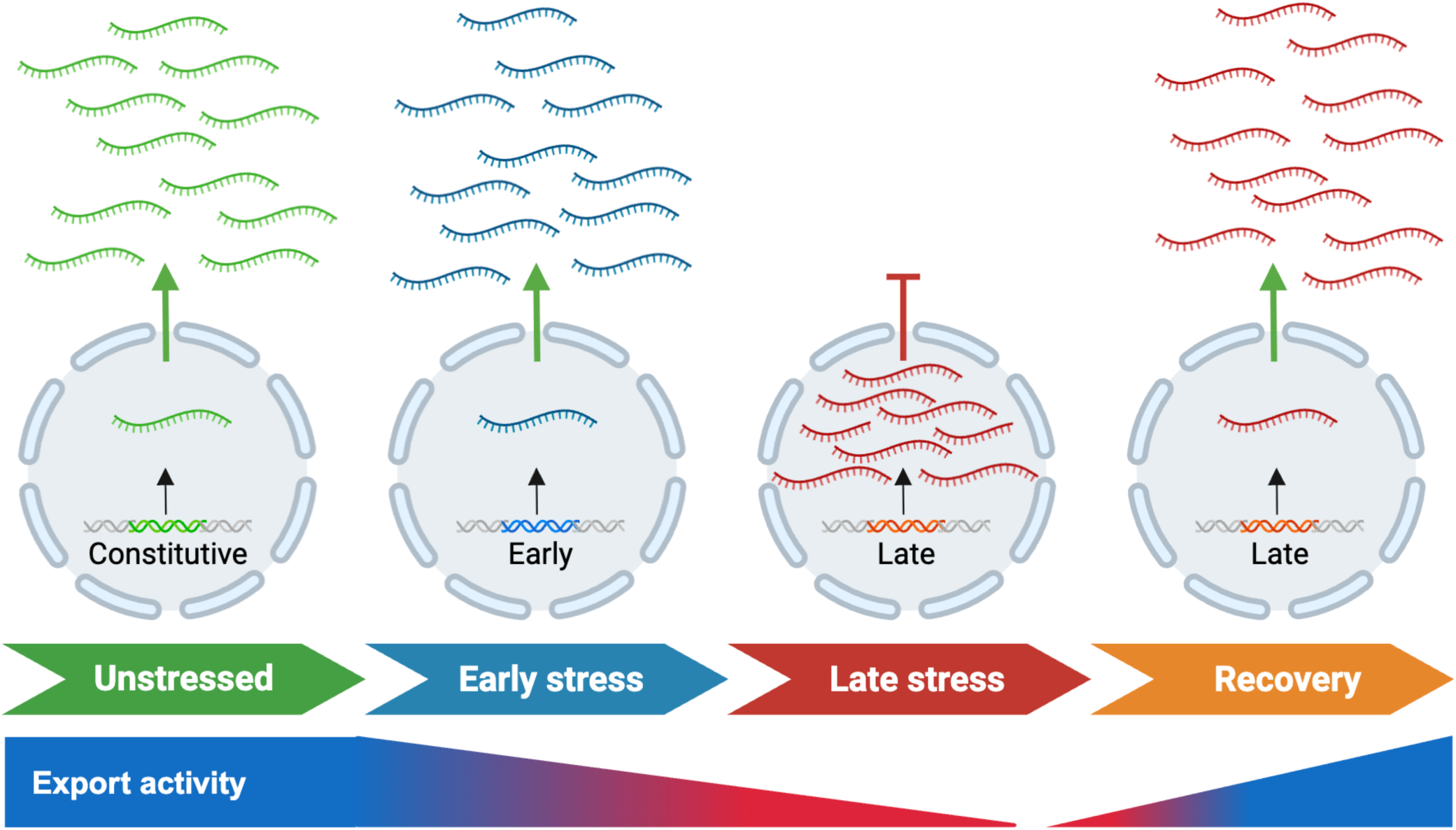
Model: temporal gating dictates stress-induced mRNA export from the nucleus. In unstressed conditions, constitutive mRNAs are generally exported to the cytoplasm after transcription. During arsenite and heat stress, early transcribed mRNAs are exported from the nucleus. The progressive inhibition of export activity results in nuclear retention of late-induced mRNAs during stress. Upon recovery from stress, mRNA export is renewed and nuclear-retained transcripts can localize to the cytoplasm.

The model that stress-induced transcripts are selectively licensed for export from the nucleus is largely based on observations in baker’s and fission yeast that show that steady state poly(A)+ RNA levels are increased in the nucleus during heat and other stresses, while HSP mRNAs are present in the cytoplasm under the same conditions (Saavedra et al. 1996; Tani et al. 1995). The selective nuclear export of HSP-encoding mRNAs in yeast is proposed to occur in a promoter- or mRNA sequence- specific manner (Saavedra et al. 1996) through the direct binding of nuclear export factors (Zander et al. 2016) and/or antisense RNAs (Coban et al. 2024) to sequence elements. However, past research did not define the contributions of transcription, nuclear export, and cytoplasmic decay to RNA localization, leaving the mechanistic basis of selective mRNA export unsubstantiated. Further, experiments in human cells reporting that heat stress causes poly(A)+ RNA to accumulate in the nucleus, while HSP70 mRNAs are exported (Gallouzi et al. 2000, 2001) were corrected several years later, yet continue to mistakenly be cited as key evidence for selective nuclear export inhibition during stress (Seidler and Sträßer 2024; Tian et al. 2026a; Khan et al. 2023). Regardless, our work does not exclude the possibility that some stress-induced mRNAs are initially exported from the nucleus through non-canonical mechanisms. Studies in mammalian cells have suggested specialized roles for export proteins TPR, Gle1, THOC5/AlyRef, and eEF1A1 in licensing HSP70 mRNA export during heat stress, although time-resolved analyses that examine transcription and mRNA decay were limited (Skaggs et al. 2007b; Katahira and Yoneda 2009; Kendirgi et al. 2005; Vera et al. 2014). Further, we observe intronless and intron-containing mRNAs in both the cytoplasmic and nuclear-retained pools, indicating that bypassing quality control mechanisms is unlikely to explain the cytoplasmic localization of some stress-induced mRNAs. Our findings appear to be generalizable across two HSR-inducing stresses and diverse cell types, and suggest that recent studies in yeast evaluating the impacts of transcriptional timing on cytoplasmic events during stress are unlikely to extend to human cells (Zedan et al. 2025; Glauninger et al. 2025). Our results, together with recent work suggesting HSF1-induced mRNAs increase in the nucleus during heat stress (Dierks et al. 2025), demonstrate progressive nuclear export inhibition overrides specific licensing mechanisms during stress.

An outstanding question is how nuclear export is progressively and reversibly inhibited during stress. We consider four possibilities here. First, the sequestration and/or mislocalization of nuclear export factors could reversibly suppress mRNA export. Impaired nucleocytoplasmic protein transport and RAN gradient disruption occurs with arsenite stress when nucleoporin and nuclear export factors localize to stress granules (Zhang et al. 2018; Vanneste et al. 2022; Hochberg-Laufer et al. 2019; Yasuda et al. 2006; Kelley and Paschal 2007; Chatterjee and Paschal 2015; Kodiha et al. 2004). While it is unclear if stress granules are required for impaired trafficking of reporter proteins (Zhang et al. 2018; Vanneste et al. 2022; Hochberg-Laufer et al. 2019), these findings suggest global defects in nucleocytoplasmic transport occur during stress.

Second, nuclear pore complexes may be damaged or modified during stress when global translation suppression would block the production of replacement proteins. In line with this possibility, ubiquitinated proteins purified from stressed cells include those that mediate nucleocytoplasmic transport (Maxwell et al. 2021), suggesting they are marked for degradation and replacement after stress. Further, previous studies have demonstrated nuclear pore modifications occur during oxidative stress that are proposed to disrupt nucleocytoplasmic transport (Kátai et al. 2016; Zachara et al. 2004; Zhang et al. 2020; Yoshimura et al. 2013; Kodiha et al. 2004). These observations suggest that global mRNA export could be inhibited due to modification or damage of nuclear pore components or export factors that are reversed or replaced during the recovery from stress.

A third possibility is that impaired quality control, processing defects, or damage could retain mRNAs in the nucleus. Heat stress can cause splicing changes (Shalgi et al. 2014), and endogenous and reporter HSP70 RNAs colocalize with nuclear speckles (Hu et al. 2010; Jolly et al. 1999). Further, the levels of poly(A)+ RNA increase in nuclear speckles with splicing factors in heat and arsenite stress (McIntyre et al. 2025). However, while we observe some instances of altered splicing in arsenite stress, retained mRNAs were diffusely localized in the nucleus, and the presence or absence of introns did not predict whether RNAs would be exported or retained. While subnuclear mRNA assemblies are observed in heat stress, arsenite stressed cells do not exhibit these assemblies and therefore they are likely not required for nuclear mRNA retention. Further, *HSPA1A* mRNAs do not harbor introns and could be exported from the nucleus early, but not late, during arsenite stress. It is also unlikely that irreversible damage to mRNAs (e.g., oxidation) is the basis of nuclear retention, as their export appears to be renewed after stress.

Fourth, mRNA modifications could contribute to nuclear RNA retention. m6A modified mRNAs increase in the nucleus during heat stress (Dierks et al. 2025). Further, increased levels of the m6A reader YTHD2 and increased m6A modifications in the 5’ UTR of transcripts including *Hspa1a* were reported during the recovery from heat stress (Zhou et al. 2015; Dierks et al. 2025). Any of these mechanisms could cause a global defect in nuclear export, which would explain the widespread, general nuclear retention of stress-induced and reporter mRNAs during stress that we observed. Future work will interrogate these potential mechanisms to clarify how nuclear export is reversibly inhibited during stress.

Regulated nuclear export could play a major role in gene reprogramming during and after stress. Our findings indicate a global, progressive inhibition of nuclear export during heat and arsenite stress from which stress-induced mRNAs are not exempt. Thus, the majority of stress-induced transcripts are inhibited from interacting with the translation machinery in the cytoplasm. Nuclear mRNA retention may safeguard the transcriptome and nascent proteome from oxidative stress and other damaging effects of stress that predominantly occur in the cytoplasm. The nuclear-retained mRNAs are longer on average, and would be predicted to be more susceptible to stress-induced damage during translation. An additional explanation is that these retained mRNAs are translationally buffered to delay their expression until stress recovery. Indeed, HSP70 protein expression can negatively regulate the HSR; thus partial nuclear retention of *HSPA1A/B* could prolong HSF1-mediated transcription until *HSPA1A* is fully exported and stress is resolved. In contrast, mRNAs that are rapidly exported (e.g., the antioxidant enzyme *HMOX1*) could be prioritized for translation during stress to directly mitigate stress effects such as reactive oxygen species. Thus, temporal gating of stress-induced mRNA export from the nucleus could contribute to the successive changes in gene expression that underlie the HSR and ISR in mammalian cells (Pakos-Zebrucka et al. 2016; Labbé et al. 2024; Han et al. 2013; Novoa et al. 2003; Pessa et al. 2024; Shi et al. 1998; Baler et al. 1992; Morimoto 1998). Future work will address these possibilities and explore the functional impacts of nuclear mRNA retention on cellular stress resilience.

Nuclear mRNA retention as a mechanism of gene regulation likely extends beyond the arsenite and heat stress conditions we examined. Prior work demonstrated increased RNA abundance in the nucleus suggestive of nuclear mRNA retention during stress conditions including glucose deprivation, ethanol, salt, and heat in yeast, and treatment with tubercidin or heat stress in human cells (Saavedra et al. 1996; Tani et al. 1995; Hochberg-Laufer et al. 2019; Coban et al. 2024; Zander et al. 2016; Heinrich et al. 2024; Dierks et al. 2025). Beyond the stress response, regulated nuclear export may explain the differential levels of nuclear and cytoplasmic RNA observed in certain cell types and tissues (Fazal et al. 2019; Bakken et al. 2018; Ren et al. 2023; Zaghlool et al. 2021; D’Sa et al. 2023; Bahar Halpern et al. 2015; Battich et al. 2015). For example, tissue-specific regulation of mRNA export in response to unknown biological cues may account for the ∼30% and ∼13% of mRNAs enriched in the nucleus in beta cells and liver cells, respectively, *in vivo* (Bahar Halpern et al. 2015). The accumulation of mRNAs in the nucleus after they are co- and post-transcriptionally processed would enable a more rapid response to changing conditions. Therefore, dynamic changes in global nuclear mRNA export may contribute to gene expression reprogramming in many biological contexts.

In addition to clarifying mechanisms of gene regulation, elucidating how nucleocytoplasmic export of mRNPs is regulated will be important to determine how defects in nucleocytoplasmic transport contribute to pathological contexts including viral infection, cancer, neurodegeneration, and genetic diseases (Moore et al. 2020; Martins et al. 2020; García-Aguirre et al. 2019; Coyne et al. 2021; Ziff et al. 2023). Viral non-structural proteins and RNAse L activation cause the accumulation of RNAs including those encoding viral or innate immune factors in the nucleus (Kumar and Glaunsinger 2010; Burke et al. 2022, 2021a, 2021b; Zhang et al. 2021). Differential expression, variants, and fusion proteins of nucleoporins are observed in cancers such as acute myeloid leukemia (reviewed in (Nofrini et al. 2016)). Nucleocytoplasmic transport defects and stresses are hallmarks of aging and aging-associated degenerative diseases (reviewed in (Moore et al. 2020; Cristi et al. 2023)). Nucleoporin mislocalization associated with impaired nucleocytoplasmic protein transport is observed in brain tissue from Huntington’s disease patients and model systems (Grima et al. 2017).

Additionally, postmortem motor cortex samples and iPSC-derived neurons from ALS/FTD patients exhibit reduced levels of nucleoporins and their limited localization to nuclear pore complexes (Coyne et al. 2020, 2021). Further, variants in the nucleoporin *NUP50* are associated with ALS/FTD (Megat et al. 2023) and mutations in the *GLE1* RNA export factor are associated with ALS (Kaneb et al. 2015). While the field has largely focused on dysregulation of protein nucleocytoplasmic transport in ALS/FTD and other degenerative diseases, one implication of our work is that nuclear export of mRNA may also be impaired. Indeed, altered nuclear:cytoplasmic RNA ratios are observed in models of degenerative disease associated with VCP mutations (Ziff et al. 2023). Finally, various genetic developmental diseases are associated with alleles of nuclear export factor genes including *GLE1* (Nousiainen et al. 2008), and members of the Transcription-Export (TREX) complex (Bhattacharjee et al. 2025): *THOC2* (Bhattacharjee et al. 2024), *THOC6* (Casey et al. 2016; Kumar et al. 2015; Werren et al. 2024), and *DDX39B* (Booth et al. 2025). Intriguingly, variants of *NUP93*, *NUP205*, and *XPO5* (Braun et al. 2016) are associated with nephrotic syndrome and NUP93 mutant models exhibit mRNA accumulation in the nucleus (Lee et al. 2026). An important future direction will be to interrogate if and how temporal gating of nuclear mRNA export during stress contributes to pathogenesis in disease contexts.

## Methods

### Cell culture and treatments

U-2 OS cells (female osteosarcoma) purchased from ATCC were maintained in Dulbecco’s modified Eagle medium (DMEM; Fisher Scientific) with 9% FetalGro EX (Rocky Mountain Biologicals), 1% penicillin/streptomycin (Gibco), and 1% Glutamax (Gibco) at 37 °C and 5% CO^2^. The dox-inducible *GADD34-Luc* U-2 OS cell line was described previously (Helton et al. 2025) and maintained in DMEM with 9% Fetal Bovine Serum, Tet system approved (Gibco), 1% penicillin/streptomycin, and 1% Glutamax at 37 °C and 5% CO^2^. hTERT-RPE cells were a kind gift from Dr. Kristen Verhey and were maintained in DMEM/F12 with 9% FetalGro EX, 1% penicillin/streptomycin, and 1% Glutamax at 37 °C and 5% CO^2^. iPSC-derived human spinal motor neuron (hSMNs) progenitors were acquired from BrainXell and matured for 5 days, following the manufacturer’s procedure, before stress treatment. Maturation of hSMNs was verified by morphological assessment and staining for MAP2 and FOXP1. Cells were periodically confirmed negative for mycoplasma by Hoechst staining and authenticated by morphological assessment. Cells were treated with the following chemicals and concentrations: sodium arsenite (Ricca Chemicals; 250 µM), actinomycin D (5 µg/mL or equivalent volume of DMSO at 0.1%), or heat stress (43°C in a 5% CO^2^ incubator) in complete growth medium for the indicated times. For recovery experiments, cells were washed 3 times with complete growth medium for 60 or 120 minutes.

All U-2 OS cell experiments were done using endogenously tagged eGFP-G3BP1 cells. This cell line was generated using CRISPR/Cas9 with a sgRNA (UUGCUUUGGUCAAUUCAACC; designed and validated by The Allen Institute) targeting the N-terminus of the endogenous G3BP1 gene. The sgRNA was ordered from Synthego with modified 2’-O-methyl analog first base and last 3 bases and 3’ phosphorothioate between the first 3 and last 2 bases. Wild-type U-2 OS cells were nucleofected with recombinant spCas9-2XNLS (Synthego), sgRNA, and an HDR template plasmid (Addgene #193920) and nucleofected using a Lonsa SE kit according to manufacturer protocols. GFP+ cells were pooled using a WOLF G2 cell sorter, and the pooled knock-in population was validated by western blotting and stress granule assays.

### FISH and smFISH

smFISH was done as previously described (Dunagin et al. 2015; Helton et al. 2025) and according to Stellaris protocols. Probes for *HMOX1, FOS, ZFP36, DNAJB1, DNAJA4,* and *DNAJB6* were designed using Stellaris Probe Designer (version 4.2) with a masking level of 5, an oligo length of 20, and a minimum spacing length of 2 nt. Probes for *GAPDH*, *GADD34, JUN,* and *Rluc* were described previously in (Helton et al. 2025) and *HSPA1A* and *HSPA1B* in (Moon et al. 2020). All probe sets were labeled with either ddUTP-ATTO-633 or ddUTP-ATTO-565 (Jena Bioscience) using terminal deoxytransferase (Thermo Scientific). Labeling reactions were performed for 16-24 hours at 37°C in a thermocycler and isolated using the oligo clean and concentrate kit (Zymo) following the manufacturer’s instructions and adjusted to 12.5 µM. The degree of labeling was determined by calculating DNA and fluorophore absorbance using a DeNovix DS-11^+^ spectrophotometer and probes were used if >80% of the probe set was labeled. All probe sequences can be found in **Supplementary Table S2**. Poly(A)+ RNA FISH was done with Cy5-oligo(dT) probes (IDT) and NEAT1-Quasar570 probes were purchased from Stellaris (Human NEAT1 middle, #SMF-2037-1).

All smFISH/FISH experiments were performed in 96-well or 8-well glass-bottom dishes (#1.5H). Following treatments, cells were washed once with PBS and fixed in 4% paraformaldehyde in PBS for 10 minutes. Cells were permeabilized in 0.1% Triton X-100 in PBS with Ribolock RNase inhibitor (Thermo Fisher Scientific) for 5 minutes and washed with wash buffer A (10% formamide in 2X SSC). Cells were incubated with each probe (1:75 for single mRNAs, 1:400 for Poly(A)+ RNA) in hybridization buffer (10% w/v dextran sulfate, 10% formamide in 2X SSC) for 16-24 hours at 37°C protected from light. Following hybridization cells were washed at 37°C once with wash buffer A containing NucBlue stain (Thermo Fisher Scientific) for 30 minutes, followed by a 30-minute incubation at 37°C in wash buffer A. Samples were imaged in 2X SSC using an iLas2 ring-TIRF (Gataca Systems) Nikon Ti2-E microscope with a 100X oil immersion objective (1.45 NA) and either an Andor Life 888 EMCCD or ORCA-Fusion BT Digital CMOS camera in HILO. Images were acquired in Z-stacks (13, 200 nm increments) and laser intensity, exposure time, and gain (for EMCCD) were kept constant for each channel across conditions of an experimental replicate. At least 3 frames were taken for every condition of every experiment. Representative images are max intensity projections deconvolved in Nikon NIS Elements with the Richardson-Lucy deconvolution algorithm, with brightness and contrast adjusted for clarity in ImageJ/FIJI.

### Nuclear/Cytoplasmic Fractionation, RT-qPCR, and RNA-sequencing

Nuclear/cytoplasmic fractionation was done as previously described (Bhatt et al. 2012; Ji et al. 2021; Ietswaart et al. 2024) with minor modifications. Briefly, unstressed or stressed (250 µM Arsenite; 105 minutes) cells were washed twice with ice-cold PBS and scraped into cytoplasmic lysis buffer (10 mM Tris-HCl pH 7.5, 150 mM NaCl, 0.1% NP-40 substitute, and Ribolock RNase inhibitor). Lysates were kept on ice for 5 minutes and 10% was kept aside for Total RNA isolation. The remaining lysates were overlaid on a sucrose solution (25% sucrose, 150 mM NaCl, 10 mM Tris-HCl) and nuclei were pelleted by centrifugation at 12,000xg for 10 minutes at 4°C. The supernatant was removed and respun once to remove residual nuclei. The nuclei pellet was washed with nuclei wash buffer (0.1% Triton-X 100, 0.2 mM EDTA, in PBS) and centrifuged at 7,500xg for 5 minutes at 4°C. Nuclei were resuspended in nuclei lysis buffer (300 mM NaCl, 1% NP-40 substitute, 1 mM DTT, 10 mM Tris-HCl pH 7.5, 0.2 mM EDTA, Ribolock RNase inhibitor) and incubated on ice for 10 minutes. Total, cytoplasmic, and nuclear fractions were mixed 1:3 with RNA-lysis buffer from the Quick-RNA Miniprep kit (Zymo Research) and RNA was isolated following the manufacturer’s instructions. Equal quantities of RNA were reverse-transcribed to cDNA using LunaScript RT SuperMix Kit and RT-qPCR was done with Luna Universal qPCR Master mix following the manufacturer’s instructions. qPCR was done for *GAPDH* and *MALAT1* (primers in **Supplementary Table S3**) on an Azure Cielo qPCR machine to validate nuclear/cytoplasmic fractionation before sequencing submission. Purified RNA was sent to Biostate.ai for rRNA-depleted RNA-sequencing with Barcode-Integrated Reverse Transcription and Probes for Excess RNA depletion at a depth of 20 million reads.

### Actinomycin D Assays

U-2 OS cells were stressed with arsenite (250 µM) for 45 or 105 min, or with heat stress (43°C) for 105 minutes prior to co-treatment with DMSO (0.1%) or actinomycin D (5 µg/mL) for 15 (*HSPA1A*) or 30-minute (all other RNAs tested) intervals. Parallel treatments were done using DMSO or actinomycin D for the entire time course to ensure transcriptional inhibition. smFISH/FISH and imaging was done as described above.

### Arsenite RNA-seq time course

U-2 OS cells were unstressed or treated with arsenite (250 µM) for 15, 30, 45, 75, or 105 min, washed once with PBS, and scraped into DNA-RNA shield (Zymo Research, #R1100-50) for submission to Plasmidsaurus for PolyA-Enriched RNA-sequencing at a depth of 20 million reads.

### Dox-inducible reporter experiments

For all dox-inducible experiments, cells were treated with 1 µg/mL of doxycycline (Fisher Scientific), and smFISH was performed as described above. For unstressed time-course experiments, cells were treated with dox in 15-min intervals for 60 min before fixation. To calculate the percent of cells with smFISH spots, a 5X5 scanning wizard image was taken and the number of cells positive for smFISH spots and the total number of cells were counted using ImageJ/FIJI Cell Counter plugin (Schindelin et al. 2012). For initial stress experiments and stress recovery experiments (**Figures 6C&D**), cells were treated with dox for a total of 60 min in each condition. The unstressed conditions were 60 min of dox in complete growth medium. To maintain a consistent 60-min dox-induction window across stress conditions, dox was added at staggered intervals. For actinomycin D treatments, cells were treated with dox 60 min before DMSO or actinomycin D treatment. Actinomycin D assays were done as described above. RNA half-life was calculated for the unstressed, actinomycin D treated sample. Half-life was determined by fitting the normalized relative abundance to an exponential decay function (y = e^-kt^) using the nls base function in R (V4.4.2). The half-life was calculated as the t^1/2^ = ln(2)/k for each biological replicate and averaged to determine the mean decay rate.

### Bioinformatic Analyses

#### Image Analysis

For all smFISH (excluding *NEAT1*), mRNA abundance and subcellular localization in U-2 OS and RPE cells were assessed using non-deconvolved raw images processed via the BigFISH/FISH-Quant (v0.6.2; (Imbert et al. 2022)) pipeline as previously described (Helton et al. 2025). For each of the three independent replicates (n = 3), total, nuclear, and cytoplasmic RNA counts were quantified from individual cells across three frames. Individual cell counts can be found in source data and 27-132 cells were counted for each condition as indicated in figure legends. Nuclei were segmented using a pretrained U-net model with TensorFlow (v2.3.0; (Abadi et al. 2016)) and cell segmentation was done using a watershed method on the GFP-G3BP1 channel. Proper segmentation was validated by eye for all images. Thresholds for smFISH spots were calculated for each experimental replicate and kept constant across conditions. *NEAT1* and Poly(A)+ RNA levels were determined using total intensity values calculated for individual cells and nuclei after segmentation using skimage (v0.17.2; (van der Walt et al. 2014)) regionprops in python (v3.6). smFISH foci were counted in ImageJ/FIJI iPSC-derived hSMNs using the Cell Counter plugin (Schindelin et al. 2012). All counts from individual cells are provided in source data (**Supplemental Table S4)**. The mean +/- s.e.m. is reported for each independent experiment with individual data points representing the mean of each independent replicate (diamonds).

#### RNA-seq preprocessing and alignment

For nuclear/cytoplasmic fractionation RNA-sequencing, preprocessing and alignment was done by Biostate.ai as follows. Briefly, sequencing adapters and low-quality reads were removed using Cutadapt (v4.2; (Martin 2011)). Trimmed reads were aligned to the GRCh38 human reference genome using HISAT2 (v2.2.1; (Kim et al. 2019)) with default parameters. Alignment quality and read distribution across genomic features (rRNA, intronic, intergenic, and exonic regions) were assessed via MultiQC (v1.14; (Ewels et al. 2016)) and Picard CollectRNASeqMetrics. Gene-level quantification was performed using featureCounts (v2.0.3; (Liao et al. 2014)) with GENCODE v46 annotation. Counts tables were used for downstream analysis.

For the arsenite time-course RNA-sequencing, preprocessing and alignment was done by Plasmidsaurus as follows. Read quality control and adapter trimming were performed using FastP (v0.24.0; (Chen et al. 2018)), including poly-X tail trimming, 3’ quality-based trimming (minimum Phred score of 15), and a minimum length requirement of 50 bps. Trimmed reads were aligned to GRCh38 human reference genome using the STAR aligner (v2.7.11; (Dobin et al. 2013)) with non-canonical splice junction removal. To account for library amplification biases, PCR and optical duplicates were removed based on unique Molecular Identifiers using UMICollapse (v1.1.0; (Liu 2019)). Gene-level expression was quantified using featureCounts (Subread package v2.1.1; (Liao et al. 2014)) with strand-specific counting and fractional assignment for multi-mapping reads. Reads were assigned to genomic features and grouped by gene ID using the GENCODE v46 annotation. Counts tables were used for downstream analysis.

#### Differential expression analysis

Differential expression analysis was performed using the DESeq2 package (v1.46.0; (Love et al. 2014)) in R (v4.4.2). Raw counts were filtered to remove lowly expressed genes, requiring a minimum of 10 reads in at least three samples. To account for high dispersion of fold change estimates for genes with low counts, log^2^ fold change (LFC) shrinkage was applied using the apeglm algorithm (Zhu et al. 2019). Significance was defined by using an adjusted p-value (Benjamini-Hochberg) of < 0.05 and a LFC threshold ≥ 0.585 (representing a 1.5-fold change). Visualization was performed using the EnhancedVolcano (v1.24.0; (Blighe et al. 2018)) and ggplot2 (v4.0.2; (Wickham 2016)) packages. Temporal classification (**Figure 5**) was restricted to protein-coding genes: ’early’ genes required significance at the 45-minute mark (vs. 0 min), while ’late’ genes were defined by significant upregulation from 45 to 105 minutes. In cases where a gene met both criteria, it was assigned to the early category.

#### Nuclear/Cytoplasmic Enrichment

Subcellular transcript distribution was quantified by calculating nuclear and cytoplasmic localization for each gene, defined by the log^2^ ratio of nuclear to cytoplasmic normalized counts. To generate these values, count matrices were filtered to exclude low-abundance genes (removal of genes with < 10 reads) prior to Median of Ratios normalization of nuclear and cytoplasmic reads. A pseudocount of 1 was added to the mean normalized counts for each fraction prior to ratio calculation to avoid division by 0. To prioritize stress-responsive transcripts with coding potential, the analysis was restricted to protein-coding genes significantly upregulated in the total RNA-seq fraction (LFC ≥ 0.585, p < 0.05). This filtering strategy was employed to minimize transcriptional noise originating from pseudogenes or non-coding RNA species. *MALAT1*, *NEAT1*, and *GAPDH* were kept as nuclear and cytoplasmic gene fiduciaries. Genes were classified as Nuclear Enriched (Log2(Nuc/Cyto) ≥ 0.585), Cytoplasmic Enriched (Log2(Nuc/Cyto) ≤ -0.585), or Intermediate.

#### Transcript Features Analysis

Gene length, transcript length, and intron number were retrieved from Ensembl via biomaRt using the Ensembl Release 115 (September 2025) database for the GRCh38.p14 human reference genome. To avoid redundancy in calculations by alternative transcript isoforms, the principal transcript was identified for each gene using the following selection priority: MANE Select status, followed by Ensembl Canonical status, and finally the longest transcript length. Transcript length and exon counts were extracted for these representative isoforms; intron counts were subsequently calculated as exon count - 1. Over representation of transcription factor motifs was performed using gprofiler2 (v0.2.4;(Raudvere et al. 2019)) in R with significance determined using Benjamini-Hochberg FDR correction.

#### smFISH/RNA-seq correlation analysis

To correlate RNA-seq and smFISH measurements, the nuclear percentage of each transcript was calculated as the fraction of normalized nuclear reads over the sum of nuclear and cytoplasmic reads. These values were plotted against smFISH nuclear percentages using the DMSO plus arsenite “time 0” condition (equivalent to 105 min arsenite) for *HSPA1A, HMOX1, PPP1R15A, HSPA1B, DNAJB1, FOS, DNAJB6, DNAJA4,* Poly(A)+ RNA, and *JUN* (**Figures 3 and 4**), and the 105 min arsenite timepoint for *NEAT1* and *GAPDH* (**Figure 1**). Linear regression and plotting were performed in R (v4.4.2) using the base lm() function.

#### Trajectory analyses

To monitor temporal expression changes from the arsenite RNA-seq time-course, raw transcript counts were normalized in DESeq2. Count matrices were filtered to exclude low-abundance genes (total reads < 10) prior to Median of Ratios normalization to account for differences in sequencing depth across libraries. Normalized counts were averaged across three independent replicates for each time point. Temporal expression was quantified as fold change relative to the 0-minute time point (unstressed). A pseudocount of 1 was applied to both numerator and denominator to avoid division by 0. Visualization was performed using the ComplexHeatmap package (v2.22.0; (Gu 2022)) color scaled to represent magnitude of induction. For categorization of nuclear, cytoplasmic, and intermediate trajectories, normalized counts from the time-course experiment of all genes in each respective group were averaged and plotted categorically.

#### IGV plots

To obtain IGV screenshots genes of interest, BAM files were converted BigWig files using the bamCoverage tool from deepTools (v3.5.6; (Ramírez et al. 2014)). Tracks were generated with a bin size of 1 bp and coverage was normalized using counts per million (CPM). Normalized bigWig files were visualized using IGV with the Human (hg38) reference and coverage was autoscaled across groups.

## Data Availability

RNA-seq files were submitted to GEO and are accessible at GSE327358. All code used for data analysis on RNA-seq datasets are available at https://github.com/noahshanehelton/Temporal-Gating-RNA-seq-Analysis. Source data is provided in **Supplementary Table S4**.

## AI Declaration Statement

Generative AI (Claude Opus 4.6) was used for language refinement in editing the manuscript.

## Acknowledgements

We thank members of the Moon lab for helpful feedback and discussions. Figure 7 was made using BioRender. This work was supported by grants to S.L.M. from the National Institutes of Health (R35GM146711) and the Chan Zuckerberg Initiative.

## Author contributions

Conceptualization - B.D., N.S.H., S.L.M.; Data curation - N.S.H.; Formal analysis - N.S.H.; Funding acquisition - S.L.M.; Investigation - N.S.H.; Methodology - N.S.H., B.D., S.L.M.; Project administration - B.D., S.L.M.; Resources - B.D., N.S.H.; Software - N.S.H.; Supervision - S.L.M.; Validation - N.S.H., S.L.M.; Visualization - N.S.H., S.L.M., B.D.; Writing - original draft - N.S.H., S.L.M.; Writing - review & editing - N.S.H., B.D., S.L.M.

