## Supplemental Figures for "Temporal gating dictates stress-induced transcript export from the nucleus"

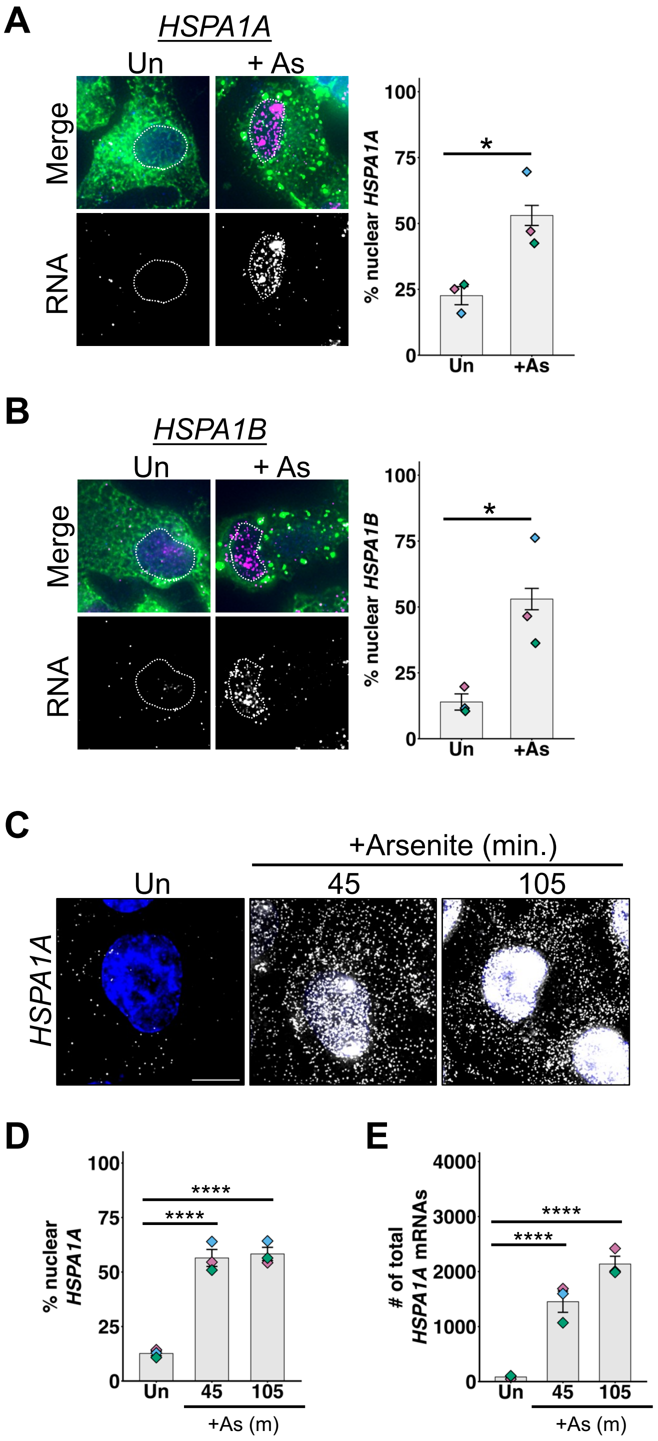


**Figure S1. HSP70 mRNAs accumulate in the nucleus during arsenite stress in human iPSC-derived spinal motor neurons and hTERT-RPE cells.** *HSPA1A* (A) and *HSPA1B* (B) (magenta) in iPSC-derived spinal motor neurons in unstressed and arsenite-stressed ("+ As", 250 µM arsenite, 45 min.) conditions.. *Left*: Representative images with immunofluorescence to G3BP1 (green) to mark the cytoplasm and FOXP1 (blue) to mark the nucleus. *Right*: quantification (average +/- s.e.m.) of percent RNA in the nucleus from n=3 independent replicates (27 cells). (C) Representative images of *HSPA1A* (white) localization in hTERT-RPE cells that are unstressed or stressed with arsenite (250 µM) for 45 or 105 min. Nuclei stained with Hoechst (blue). The percent nuclear (D) and total number of (E) *HSPA1A* mRNAs in hTERT-RPE cells as in (C) from n=3 independent experiments (70 - 81 cells counted per condition). Scale bars = 10 µm. Statistical significance between conditions was assessed by one-way ANOVA, (*) *P < 0.05,* (****) *P < 0.001.*

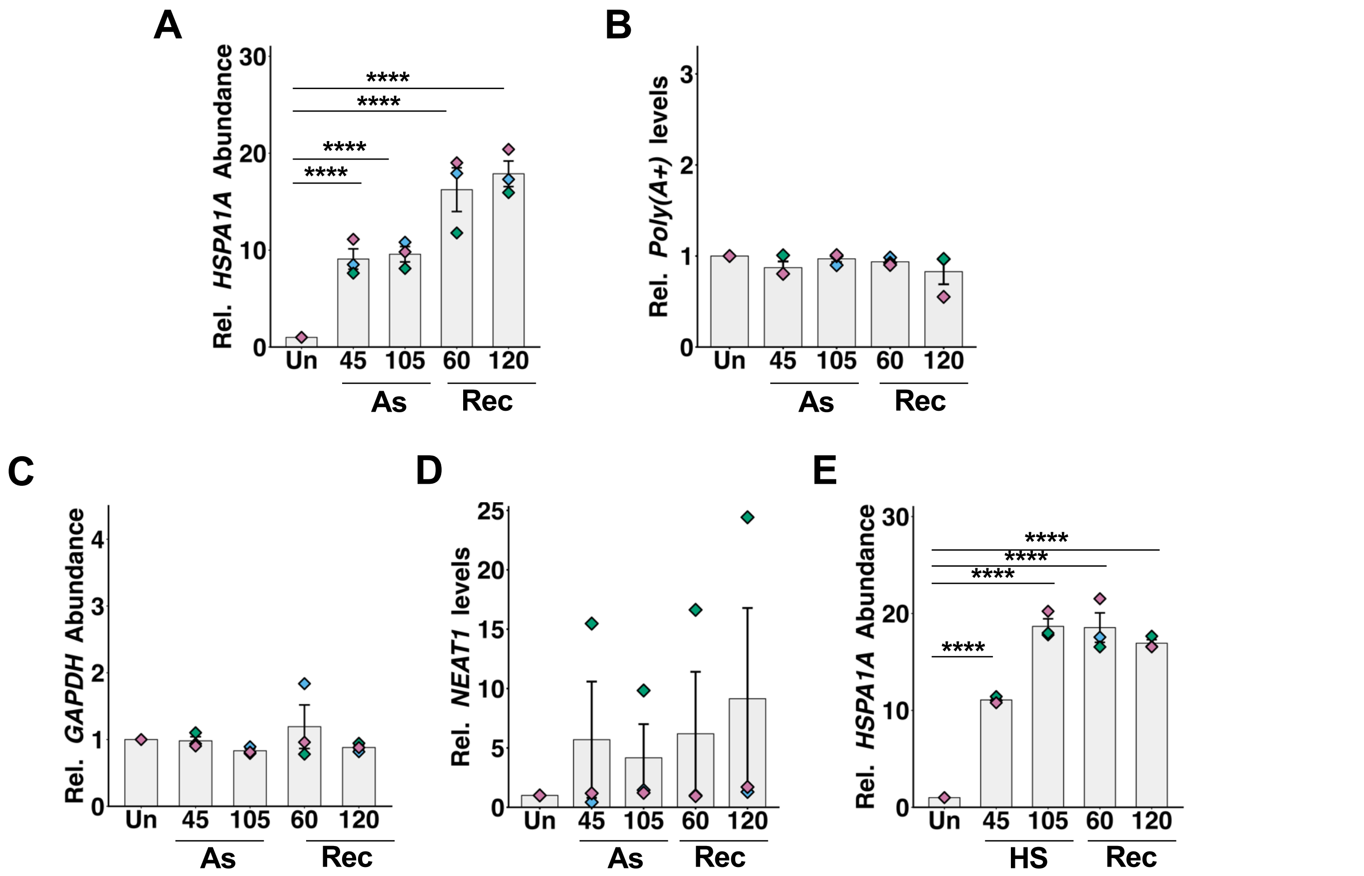


**Figure S2. Relative levels of RNAs during and after arsenite and heat stress.** Cells were unstressed ("Un") or stressed for 45 or 105 min, or stressed 45 min then recovered ("Rec") for 60 or 120 min. Average (+/- s.e.m.) relative abundance of *HSPA1A* (A), poly(A) RNA (B), *GAPDH* (C), and *NEAT1* (D) during arsenite stress ("As") from smFISH data is shown. (E) Average relative abundance (+/- s.e.m.) of *HSPA1A* during heat stress ("HS") from smFISH data is shown. Results are from n = 3 independent experiments (46-132 cells counted per condition). Statistical significance determined using one-way ANOVAs with (****) $P < 0.001$.

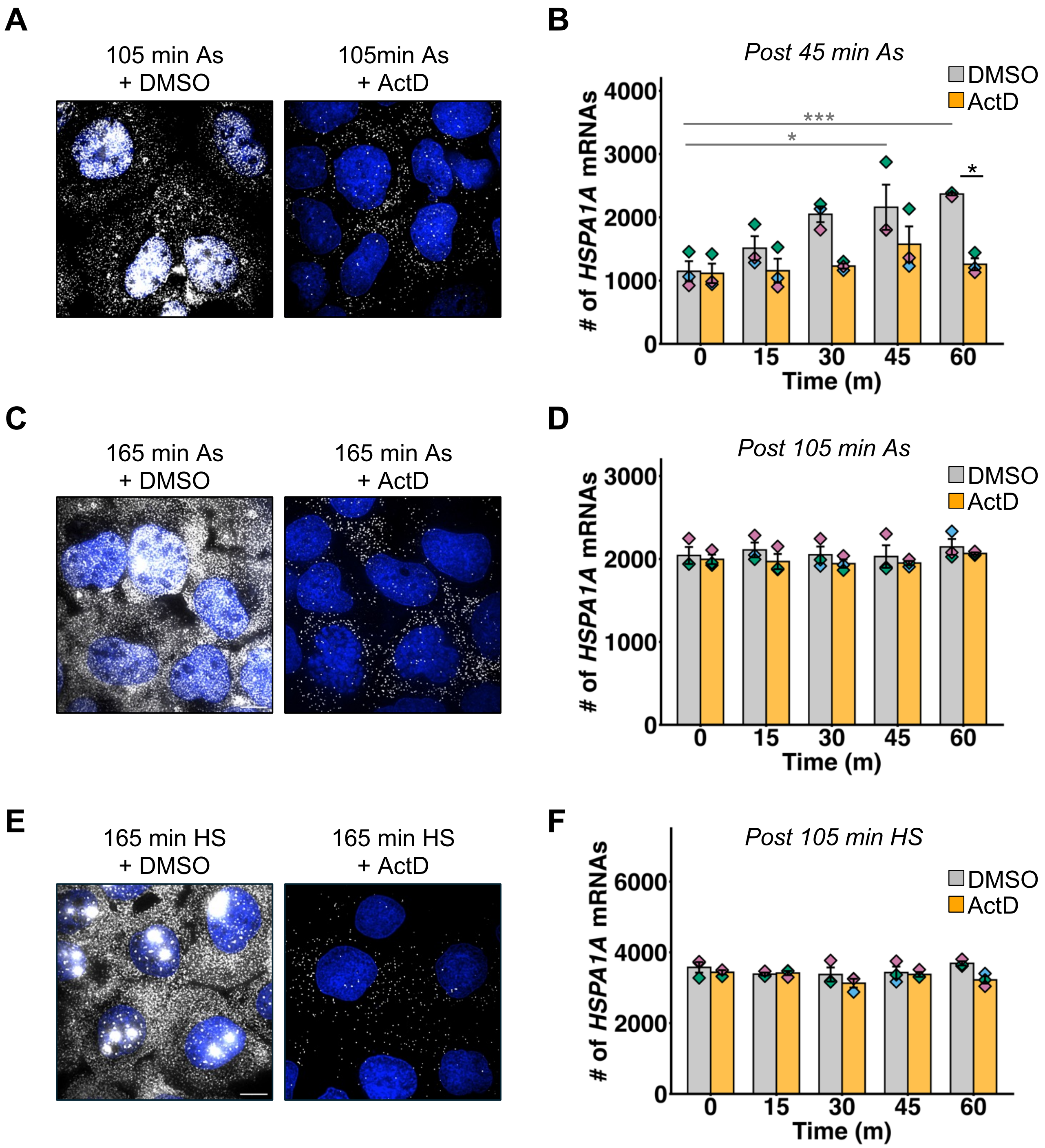


**Figure S3. Validation of actinomycin D activity and number of *HSPA1A* mRNAs per cell in each condition during arsenite and heat stress.** (A) Representative images of *HSPA1A* (white) in cells treated with carrier DMSO (0.1%) or actinomycin D ("ActD", 5 µg/mL) for 105 min during arsenite stress ("As"; 250 µM). Nuclei visualized with Hoechst (blue). (B) Average +/- s.e.m. of the number of *HSPA1A* mRNAs per cell in samples treated with DMSO or actinomycin D for 0 - 60 min. starting at 45 min of arsenite stress. (C) As in (A), for 165 min. unstressed or arsenite-stressed conditions. (D) As in (B), for the actinomycin D time course starting at 105 min of arsenite stress. (E) As in (A), for cells treated for 165 min. with DMSO or actinomycin D during heat stress ("HS"). (F) As in (D), for the actinomycin D time course starting at 105 min. heat stress (43 °C). Results represent n=3 independent experiments, with statistical significance assessed using one-way ANOVAs; grey indicates comparisons between DMSO time points, and black indicates comparisons between DMSO and ActD conditions; (*) $P < 0.05$, (***) $P < 0.005$.

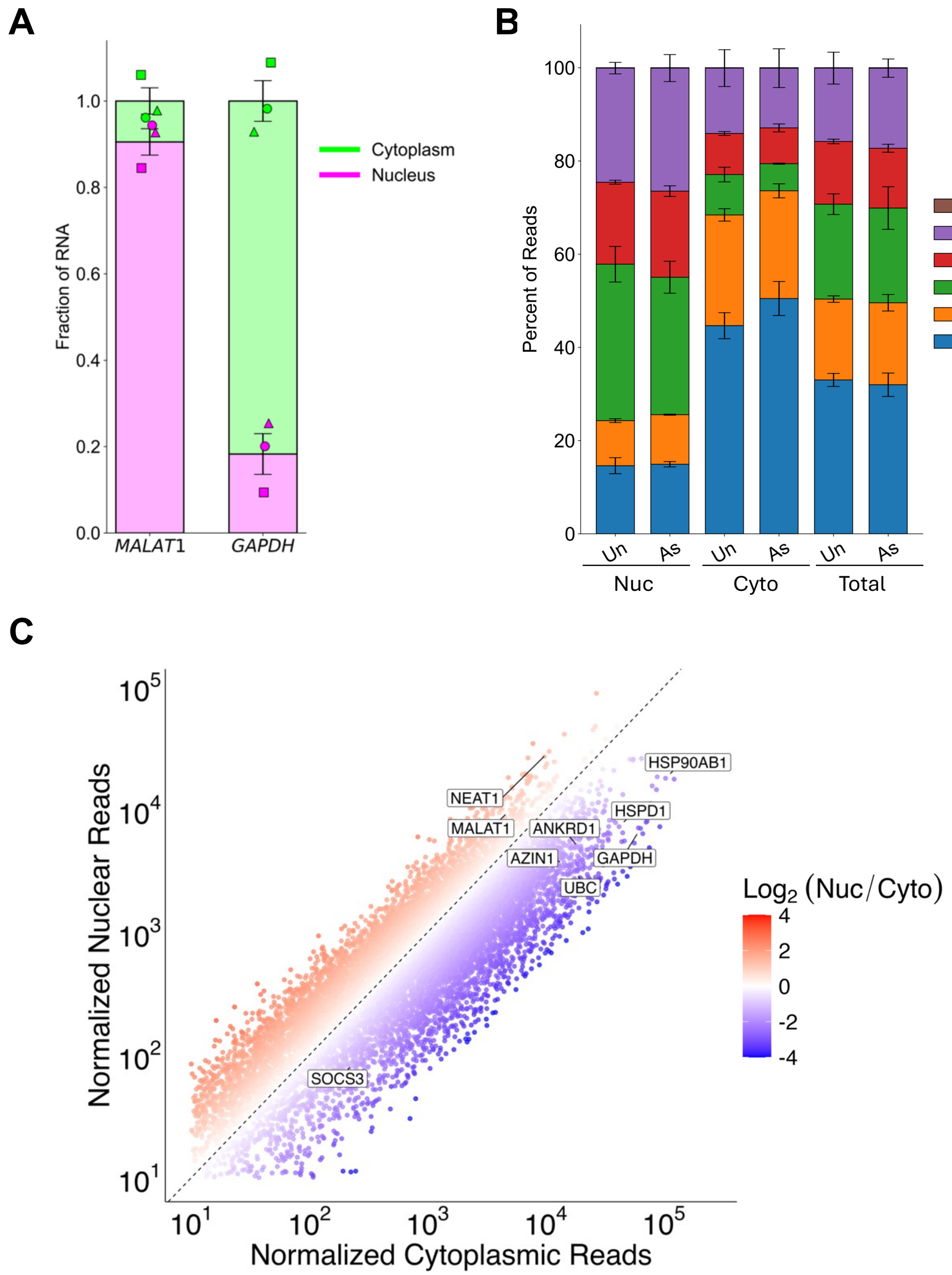


**Figure S4. Validation of nuclear:cytoplasmic fractionation.** (A) RT-qPCR analysis of *GAPDH* and *MALAT1* from nuclear and cytoplasmic fractions of unstressed cells. Stacked bars show the mean fraction +/- s.e.m. of each transcript in the nucleus (magenta) or cytoplasm (green), with individual points representing each independent replicate (n=3). (B) Stacked bar graph showing the average percentage of RNA-seq reads assigned to genomic features in nuclear (“nuc”), cytoplasmic (“cyto”), and total fractions under unstressed (“Un”) or arsenite-stressed (“As”; 250 μM, 105 min) conditions. Reads were classified using Picard CollectRnaSeqMetrics into Unaligned (brown), rRNA (purple), Intergenic (red), Intronic (green), UTR (orange, 5’ and 3’ UTRs), and Coding (blue, CDS exons only). Bars represent mean fraction percent +/- s.e.m. (n = 3 for all conditions except unstressed nuclear n = 2). (C) Comparison of normalized nuclear vs cytoplasmic protein-coding mRNA RNA-seq reads from unstressed samples (n=3 for cytoplasmic, n=2 for nuclear). $Log_2$ fold change in partitioning shown with red indicating more nuclear and blue indicating more cytoplasmic transcripts. Labeled RNAs include nuclear lncRNAs *NEAT1* and *MALAT1*, and cytoplasmic mRNA *GAPDH*. Other labeled mRNAs represent cytoplasmic-enriched late-induced transcripts that are highly cytoplasmic and abundant before stress.

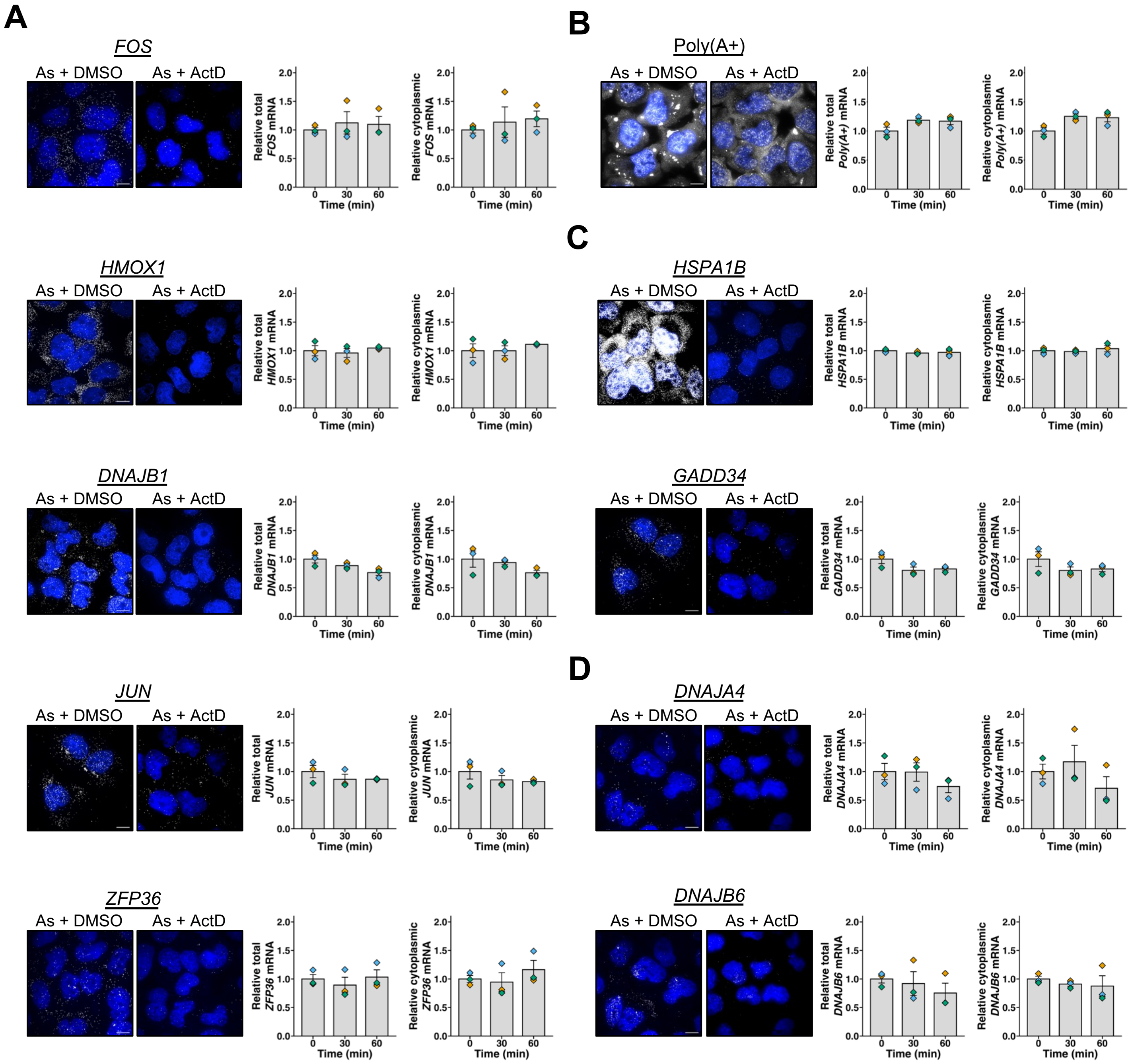


**Figure S5. Validation of actinomycin D activity and relative total and cytoplasmic mRNA abundances during actinomycin D shutoffs starting at 105 min arsenite stress.** Representative smFISH/FISH images (at left) and relative total (middle) and cytoplasmic levels (at right) of (A) Cytoplasmic, (B) poly(A)+ RNA, (C) intermediate mRNAs, and (D) nuclear-enriched mRNAs during 60-min actinomycin D ("ActD", 5 µg/mL) or carrier control DMSO (0.1%) time course experiments starting at 105 min of arsenite stress ("As"). RNA shown in white and nuclei in blue; average +/- s.e.m. from n = 3 independent replicates with each replicate average shown as diamonds; 51-119 cells counted per condition.

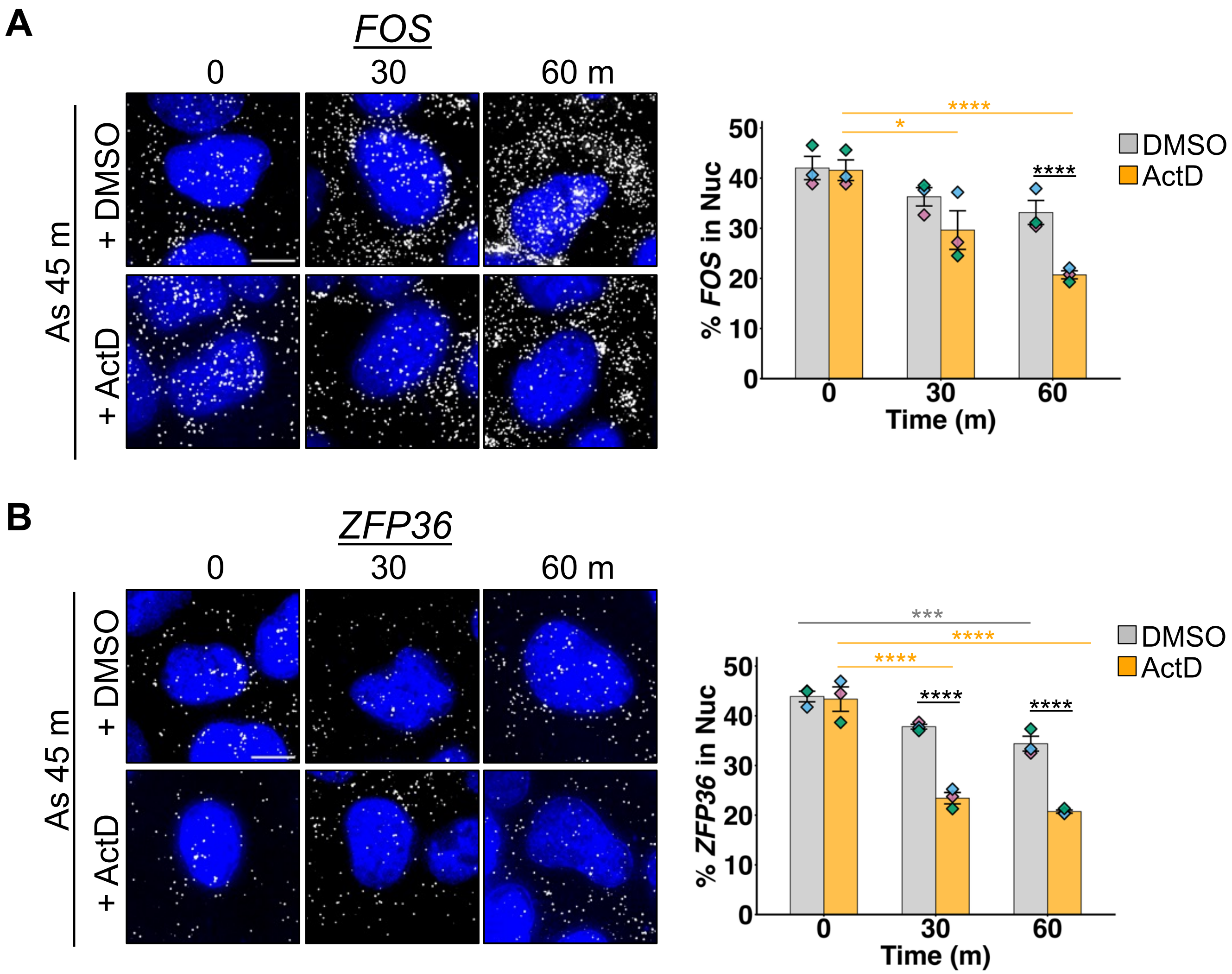


**Figure S6. Export of *FOS* and *ZFP36* starting 45 min after arsenite stress.** Cells were stressed with arsenite ("As"; 250 µM) for 45 min, then co-treated with actinomycin D ("ActD", 5 µg/mL) or DMSO (0.1%) for 60 min. *Left*: representative smFISH images. *Right*: Quantification of n = 3 independent replicates showing the mean +/- s.e.m. of the nuclear percentage of (A) *FOS* and (B) *ZFP36* over time. Grey bars indicate DMSO treatment and orange represent ActD treatment. Diamonds represent the average of each independent replicate. One-way ANOVA tests for significance were done between DMSO time points (grey), ActD time points (orange) or between DMSO and ActD time points (black) with (*) $P < 0.05$, (***) $P < 0.005$, (****) $P < 0.001$.

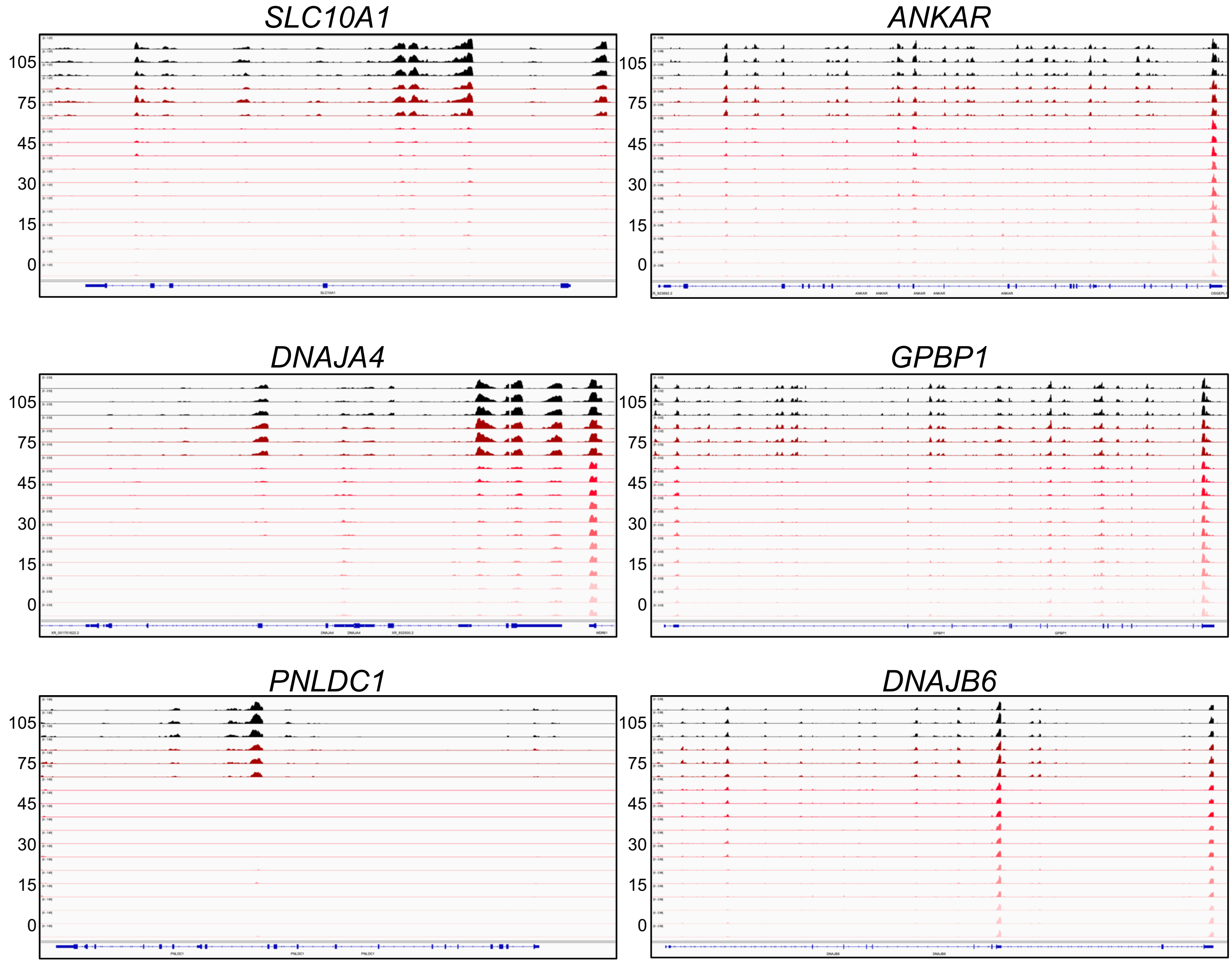


**Figure S7. Late-induced mRNA RNA-seq read coverage over time during arsenite stress.** IGV screenshots for representative late-induced transcripts. Each row represents read coverage for individual replicates (n = 3) for each time point from 0 to 105 min of arsenite stress.

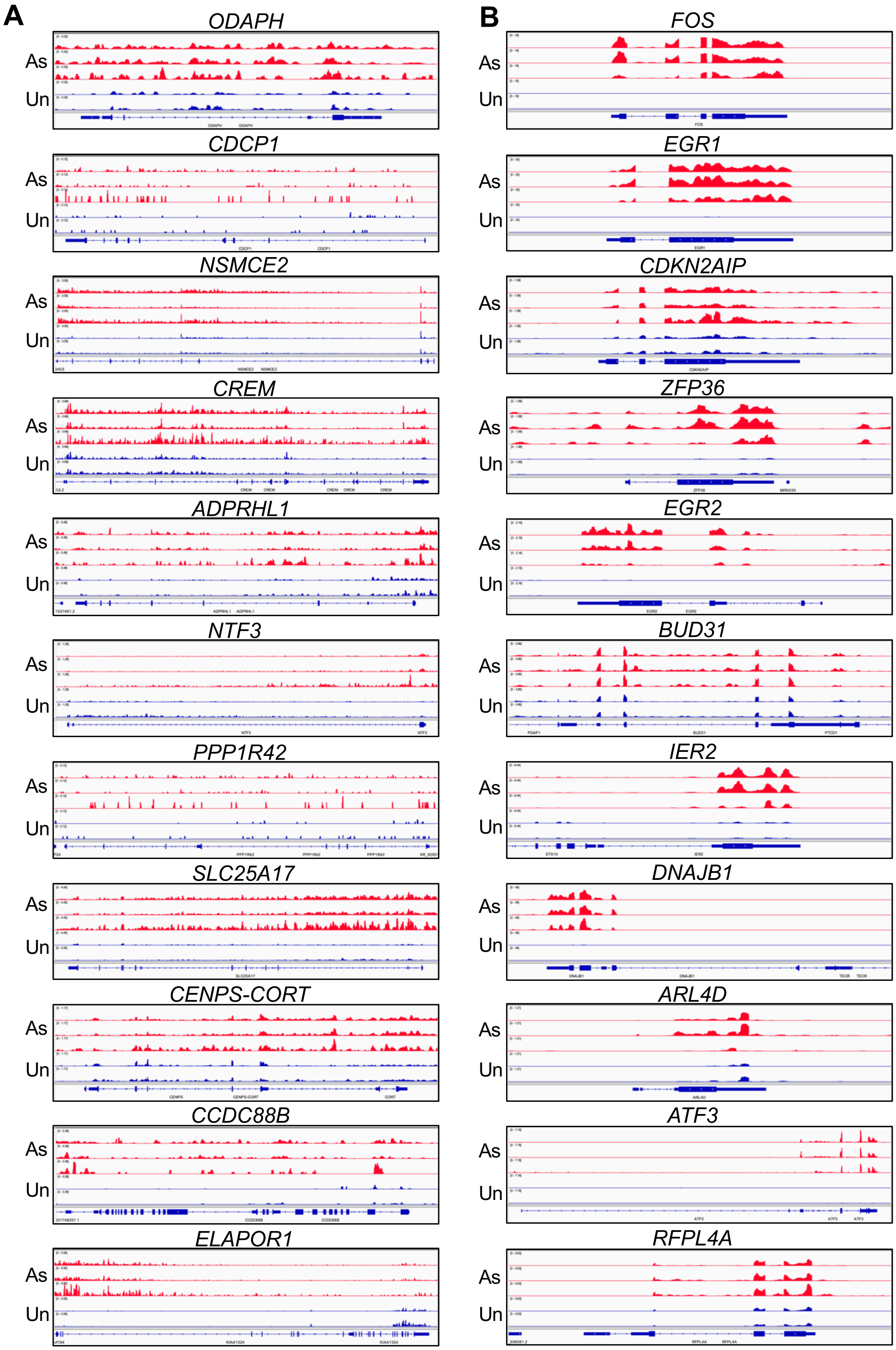


**Figure S8. Early-induced mRNA nuclear RNA-seq read coverage.** IGV screenshots of the nuclear compartment for early, nuclear genes (A) and early, cytoplasmic genes (B) in stressed (arsenite, 250 μM, 105 min) or unstressed conditions. Each row represents read coverage for individual replicates (n = 3 for arsenite, n = 2 for unstressed).

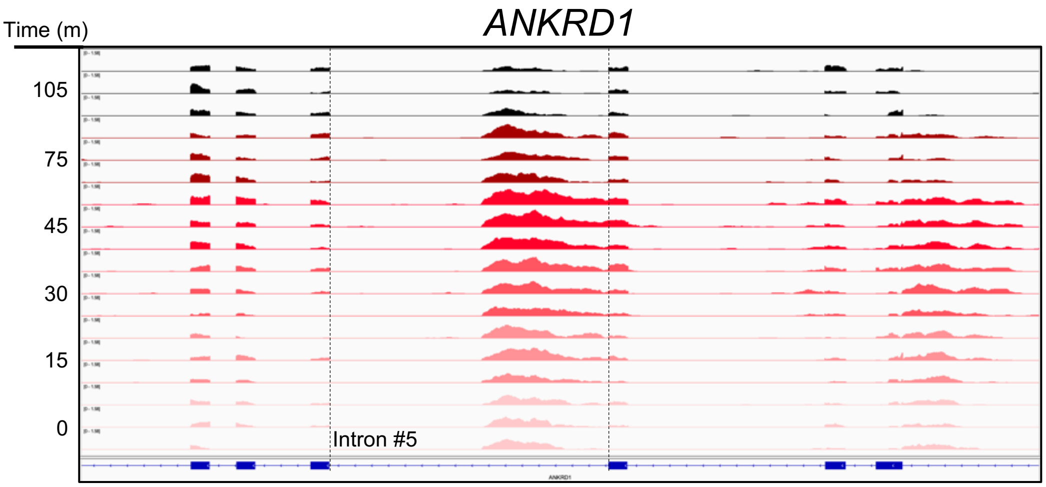


**Figure S9. *ANKRD1* intron 5 read coverage decreases over time during arsenite stress.** IGV screenshot of *ANKRD1* over an arsenite stress (250 μM) time-course. Each row represents read coverage for individual replicates (n = 3) for specific time points from 0 to 105 min of arsenite stress.

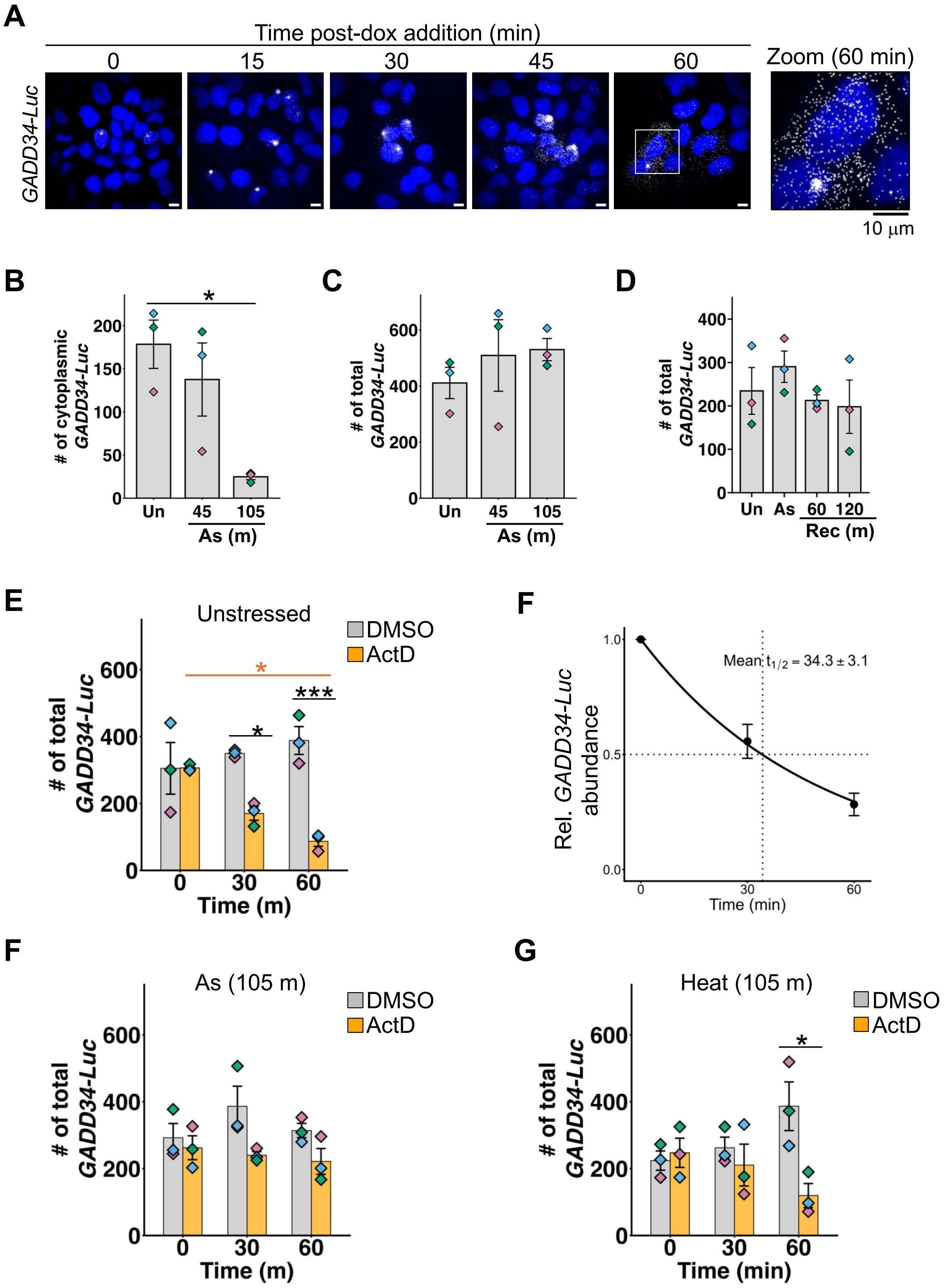


**Figure S10. Dox-inducible *GADD34-Luc* reporter mRNA induction and abundance.** (A) Representative images of dox-induced *GADD34-Luc* (white) and nuclei (blue). (B-C) Bar graphs of average +/- s.e.m. cytoplasmic (B) and total *GADD34-Luc* mRNA counts per cell from Figure 6D. (D) Average +/- s.e.m. of the total *GADD34-Luc* mRNAs per cell from unstressed DMSO (0.1%) or actinomycin D ("ActD", 5 µg/mL). (E) mRNA half-life of *GADD34-Luc* mRNA abundance in cells treated with ActD, with the fitted curve shown from n = 3 independent replicates and the average +/- s.e.m. half-life (min) shown. Average +/- s.e.m. total *GADD34-Luc* mRNAs per cell from (F) arsenite ("As", 250 µM 105 min) or (G) heat (43°C, 105 min) stressed cells co-treated with DMSO or ActD. One-way ANOVAs were done to determine statistical significance across time points; orange indicating significance between ActD timepoints and black indicating significance between DMSO and ActD time points; (*) *P < 0.05,* (***) *P < 0.005.*
