## Supplemental Table Legends for "Temporal gating dictates stress-induced transcript export from the nucleus"

**Supplementary Table S1.** Nuclear, cytoplasmic, and intermediate upregulated stress-induced genes, related to Fig. 3.

**Supplementary Table S2.** Oligonucleotide sequences for smFISH probes used to detect *HSPA1A, HSPA1B, DNAJB1, DNAJB6, DNAJA4, GADD34, JUN, FOS, ZFP36, GAPDH, HMOX1,* and *Luciferase*. Related to Fig. 1, 2, 4, 6 and Supplemental Fig. S1, S2, S3, S5, S6, and S10.

**Supplementary Table S3.** qPCR primers used to detect *GAPDH* and *MALAT1,* related to Supplemental Fig. S4.

**Supplementary Table S4.** Source data for all figures.
